# USP28 enables oncogenic transformation of respiratory cells and its inhibition potentiates molecular therapy targeting mutant EGFR, BRAF and PI3K

**DOI:** 10.1101/2021.09.06.459088

**Authors:** Cristian Prieto-Garcia, Oliver Hartmann, Michaela Reissland, Fabian Braun, Süleyman Bozkurt, Carmina Fuss, Christina Schülein-Völk, Alexander Buchberger, Marco A. Calzado Canale, Mathias Rosenfeldt, Ivan Dikic, Christian Münch, Markus E. Diefenbacher

## Abstract

Oncogenic transformation of lung epithelial cells is a multi-step process, frequently starting with the inactivation of tumor suppressors and subsequent activating mutations in proto-oncogenes, such as members of the PI3K or MAPK family. Cells undergoing transformation have to adjust to changes, such as metabolic requirements. This is achieved, in part, by modulating the protein abundance of transcription factors, which manifest these adjustments. Here, we report that the deubiquitylase USP28 enables oncogenic reprogramming by regulating the protein abundance of proto-oncogenes, such as c-JUN, c-MYC, NOTCH and ΔNP63, at early stages of malignant transformation. USP28 is increased in cancer compared to normal cells due to a feed-forward loop, driven by increased amounts of oncogenic transcription factors, such as c-MYC and c-JUN. Irrespective of oncogenic driver, interference with USP28 abundance or activity suppresses growth and survival of transformed lung cells. Furthermore, inhibition of USP28 via a small molecule inhibitor reset the proteome of transformed cells towards a ‘pre-malignant’ state, and its inhibition cooperated with clinically established compounds used to target EGFR^L858R^, BRAF^V600E^ or PI3K^H1047R^ driven tumor cells. Targeting USP28 protein abundance already at an early stage via inhibition of its activity therefore is a feasible strategy for the treatment of early stage lung tumours and the observed synergism with current standard of care inhibitors holds the potential for improved targeting of established tumors.

## Introduction

In the past decade, with the advent of targeted therapy, great advancements towards the treatment of progressed Non-Small Cell Lung Cancer (NSCLC) in distinct patient cohorts were achieved(1), while patients with early disease do not benefit from these new treatments(2, 3). For this cohort, the curative treatment, still today, is the surgical resection of a lung lobe. This is a severe procedure, inflicting major damage, requires an extended recovery time and can result in therapy induced mortality(4, 5). Furthermore, therapy failure in late stage tumours by establishment of treatment escape mechanisms is a common observation in NSCLC, significantly affecting patient survival(6, 7). Overall, survival rates have only marginally improved and most patients still succumb to the disease(1).

Therefore, targeting of common essential pathways and exploiting tumour intrinsic vulnerabilities holds the potential to not only improve current treatment for late stage, but also for early stage patients. One central cellular component tumour cells alter during oncogenic transformation is the ubiquitin proteasome system (UPS)(8, 9). The dysregulation of the UPS is a prerequisite for tumor cells to tolerate increased proliferation, metabolic changes, immune evasion and proteostatic stress management(10). All these processes are ‘hallmarks of cancer’ and therefore significantly contribute to disease progression, therapy failure and shorted survival. Therefore, cancer cells, when compared to non-transformed cells, are dependent on the ubiquitin system(11, 12). As a consequence, tumour cells develop exploitable dependencies towards the expression and abundance of discreet members of the UPS^12^.

Despite the prominent involvement of the UPS in cancer, our understanding of how tumour cells alter the UPS system very early in transformation is rather limited(12). The identification of essential and druggable key-players within this class of enzymes has the potential to hold novel therapeutic strategies. Deubiquitinating enzymes are such a therapeutically promising class of enzymes, as individual members can be targeted by small molecule inhibitors(13-15).

In this study, we report that the deubiquitylase USP28 presents a UPS enzyme, which is commonly upregulated during early stages of oncogenic transformation in lung cancer. Irrespective of oncogenic driver, tumour cells upregulate USP28, which stabilizes proto-oncogenes, such as c-MYC, c-JUN or NOTCH. Tumour cells are addicted to USP28 to allow oncogenic transformation and its inhibition via the small molecule inhibitor AZ1(16) partially reverts the oncogenic transformation. Finally, combining USP28 targeting with targeted therapy against commonly found oncogenic drivers potentiates treatment responses, at least *in cellulo*, indicating that the UPS system, exemplified by USP28, is a promising target structure for lung cancer.

## Materials and Methods

### Cell lines

Human basal bronchial epithelial BEAS-2B cells were originally transformed with SV40-large-T-antigen (Reddel et al., 1988). The cell line BEAS-2B was a kind gift of Prof. Marco A. Calzado Canales (Universidad de Córdoba, Hospital Reina Sofia, Córdoba, Spain). BEAS-2B Oncogenic cells were generated upon retroviral infection of BEAS-2B DIF with the next plasmids: EGFR (addgene number: #11011), EGFR L858R (addgene number: #11012), pBabe puro HA PIK3CA (addgene number: #12522), pBabe puro HA PIK3CA H1047R (addgene number: #12524), pBabe puro HA PIK3CA E545K (addgene number: #12525), pBabe puro HRAS G12D (HRAS G12D was cloned into pBabe puro in our lab) and pBabe puro BRAF V600E (addgene number: #15269). The plasmids EGFR and EGFR L858R were a gift from Matthew Meyerson (Greulich H, Chen TH et al. 2005). pBabe puro HA PIK3CA H1047R, HA PIK3CA E545K, HA PIK3CA were a gift from Jean Zhao (Zhao JJ, Liu Z, Wang L et al. 2005). pBabe Puro BRAF V600E was a gift from William Hah (Boehm et al Cell. 2007) For virus production HEK293-T cells were used. Cell lines used in this publication are listed in the supplementary table called: Consumables and resources.

### Tissue culture reagents and drugs

Cells were plated on Greiner dishes and incubated at 37 °C, 95 % relative humidity and 5 % CO2 in a cell incubator for optimal growth conditions. DIF BEAS-2B, oncogenic BEAS-2B and HEK-293T cells were cultured in DMEM (Gibco) supplemented with 10% fetal bovine serum (FCS)/ 1% Pen-Strep. UD cells were cultured in LHC-9 (Gibco) supplemented with 1% Pen/Strep. To cultivate UD BEAS-2B cells, the dishes were pre-coated with pre-coating solution composed by: 0.03% Collagen (in 0.1 M acetic acid), 0.01% Fibronectin and 0.001% BSA. UD cells were supplemented with 10% FCS to induce pre-oncogenic differentiation. Cells were routinely tested for mycoplasma via PCR.The reagents and drugs were dissolved in Dimethyl sulfoxide (DMSO). AZ1, Gefitinib, Buparlisib and Vemurafenib were purchased from Selleckchem. Drugs and reagents are listed in the supplementary table called: Consumables and resources.

### AAV, Retrovirus and Lentivirus production and purification

Adeno-associated viruses (AAVs) were generated and packaged in HEK293-T cells seeded in 15-cm cell culture dishes (60-70% confluence). Cells were transfected with the plasmid of interest (10 μg), pHelper (15 μg) and pAAV-DJ (10 μg) using PEI in ratio 2:1 (70 μg). After 96 hours, AAV Virus isolation from cells was performed as previously described (17). For Retrovirus production, HEK293 cells (70% confluence) were transfected with the babe plasmid of interest (15 μg), pUMVC (10 μg) and VSV-G (10 μg) using PEI (70 μg). After 96 H, the medium containing retrovirus was filtered (0.45 µM) and stored at -80°C. For Lentivirus production, HEK293 cells (70% confluence) were transfected with the plasmid of interest (15 μg), pPAX (10 μg) and pPMD2 (10 μg) using PEI (70 μg). After 96 H, the medium containing lentivirus was filtered (0.45 µM) and stored at -80°C.

### In vitro DNA transfection and infection

DNA transfection was performed exposing 60% confluence BEAS-2B cells plated in a 6-well cell culture dish to a mix of 2.5μg plasmid of interest, 200μl DMEM free serum and 5μl PEI (1:2 ratio). Upon 6h incubation at 37°C, 5% CO2 and 95% relative humidity, the medium was removed and substituted by DMEM (Gibco) supplemented with 10% FCS/ 1% Pen-Strep. For viral infection, 10 MOI (multiplicity of infection) of Retroviruses (LVs) were added to normal medium of the cells in the presence of polybrene (5μg/ml). Cells exposed to the viruses were incubated at 37°C, 5% CO2 and 95% relative humidity for 4 days. The infected cells were identified and selected by exposure to 2.5μg/ml Puromycin for 72h.

### RT-PCR

RNA was isolated with Peq GOLD Trifast (Peqlab), as indicated in the manufacturer’s instructions. RNA was reverse transcribed into cDNA using random hexanucleotide primers and M-MLV enzyme (Promega). Quantitative RT-PCR was performed with SYBR Green mix (ABgene) on the instrument “Step One Realtime Cycler”(ABgene) The RT-PCR program employed in this research is the following: 95°C for 15 min., 40x [95°C for 15 sec., 60°C for 20 sec. and 72°C for 15 sec.], 95°C for 15 sec. and 60°C for 60 sec. Relative expression was generally calculated with ΔΔCt relative quantification method. Melt curve was performed for all primers. For visualization purposes, Excel (Microsoft) and Affinity Designer were used as bioinformatic tools. Primers used for this publication are listed in the supplementary table called: Consumables and resources.

### Plasmids, sgRNA and shRNA Design

sgRNAs were designed using the CRISPR online tool: https://zlab.bio/guide-design-resources). shRNAs were designed using SPLASH-algorithm: http://splashrna.mskcc.org/) or RNAi Consortium/Broad Institute: www.broadinstitute.org/rnai-consortium/rnai-consortium-shrna-library. Oligonucleotides used in this publication are listed in the supplementary table called: Consumables and resources.

### Operetta analysis, Immunofluorescence, cell viability, Bliss synergy and GI50

Number of cells was quantified using Operetta High-Content Imaging System (PerkinElmer) (number of DAPI positive cells) or Invitrogen Countess II FL (number of cells after trypsinization) upon indicated treatments. For the Operetta High-Content Imaging System, cells were seeded in 384-well plates at equal density and exposed to indicated treatments. Then, cells were fixed using 4% PFA for 10 minutes and then, permealized using 0,5% Triton x100 in PBS for 5 minutes. For IF, primary antibodies (1/100) were incubated ON at 4°C, followed by subsequent incubation with the secondary antibody (1/300) for 1 hour at room temperature. After antibody exposure, samples were washed twice with PBS. Before quantification cells were stained with DAPI (ThermoFischer). For the quantification of dead cells, 1ug/ml PI was added to the cell medium of live cells for 20 minutes upon indicated treatments. For quantification of dead cells, 1ug/ml PI (Sigma Aldrich) was added to the cell medium of live cells for 20 minutes upon indicated treatments. For quantification of proliferative cells, samples were subjected to KI67 (Santa cruz ab: sc-23900; 1/100) staining by IF before imaging. Number of dead, proliferative and total cells were determined counting the number of positive nucleus for PI, KI67 or DAPI with the Harmony Software (Perkin Elmer). Bliss synergy was calculated using the total number of cells upon indicated treatments. For calculation of synergy, Combenefit software was previously described (Di Veroli GY et al 2016). GI50_50_ was generated using the online tool: www.aatbio.com. For Crystal violet cell viability, cells were stained with 0.5% Crystal violet and analyzed using ImageJ software (staining intensity is between 0 to 255). For visualization purposes, Excel (Microsoft) and Affinity Designer were used as bioinformatic tools. Antibodies used in this publication are listed in the supplementary table called: Consumables and resources.

### Immunoblot

Cells were lysed in RIPA lysis buffer (20 mM Tris-HCl pH 7.5, 150 mM NaCl, 1mM Na^2^EDTA, 1mM EGTA, 1% NP-40 and 1% sodium deoxycholate), supplemented with proteinase inhibitor (1/100) via sonication with Branson Sonifier 250 (duty cycle at 20% and output control set on level 2; 10 sonication / 1 minute cycles per sample). 50μg protein was boiled in 5x Laemmli buffer (312.5mM Tris-HCl pH 6.8, 500 mM DTT, 0.0001% Bromphenol blue, 10% SDS and 50% Glycerol) for 5 min and separated on 10% Tris-gels in Running buffer (1.25M Tris base, 1.25M glycine and 1% SDS). After separation, protein was transferred to Polyvinylidene difluoride membranes (Immobilon-FL) in Transfer Buffer (25mM Tris base, 192mM glycine and 20% methanol). Membrane was exposed to blocking buffer (0.1% casein, 0.2xPBS and 0.1% Tween20) for 45-60 min at room temperature (RT). Then, membranes were incubated with primary antbody (1/1000) in a buffer composed by 0.1% casein, 0.2x PBS and 0.1% Tween20) for 6h at room temperature (RT). Membrane was incubated with indicated secondary Antibody (1/10000) in a buffer composed by 0.1% casein, 0.2x PBS, 0.1% Tween20 and 0.01% SDS for 1h at RT. Membranes were recorded in Odyssey® CLx Imaging System, and analysed using Image Studio software (Licor Sciences). Antibodies used in this publication are listed in the supplementary table called: Consumables and resources.

### Ubiquitin Suicide Probe/Warhead DUB activity assays

Cells were resuspended in HR-buffer (50 mM Tris-HCl pH 7.4, 5 mM MgCl2, 250 mM Sucrose, 0,1 % NP-40), supplemented with Protease-Inhibitor. Lysis was performed by three freeze-thaw cycles. 25 µg of cell lysate were transferred to a new eppedornf tube and 3 µL of a 1:1:1 mixture of Ub-VME, Ub-VS, Ub-PA suicide-probes (UbiQ) resuspended in 50 mM NaOAc, 5 % DMSO were added to the mixture. In order to adjust the pH, 50 mM NaOH was added. Then, samples were mixed and incubated for 1 hour at 37°C shaking. After addition of Laemmli-buffer, samples were boiled for 5 min and applied to immunoblot was performed.

### Human lung cancer samples

Human samples were obtained from Pathology Department Córdoba (Spain), Pathology Department University Hospital Würzburg (Germany) and U.S. Biomax (lung microarray slides; slide LC2083). Informed consent was obtained from all patients. Experiments were in agreement to the principles set out in the WMA Declaration of Helsinki and the Department of Health and Human Services Belmont Report. Samples are approved under ethical approval license decret 439/2010 (Hospital Universitario Reina Sofía) and Ethics approval 17/01/2006 (University Hospital Würzburg).

### Analysis of human publicly available datasets

Oncoprints were generated using cBioportal online tool. Briefly, Oncoprints generates graphical representations of genomic alterations, somatic mutations, copy number alterations and mRNA expression changes. Graphical representations of somatic mutations were performed using the online tool Cbioportal. TCGA data was used for the different analysis. Correlation analysis and USP28 expression in different subtypes of ADC and SCC lung tumors were obtained using GEPIA’s software (Tang Z. et al. 2017). For GEPIA gene expression. the differential analysis was based on: “ TCGA tumors vs (TCGA normal)”, whereas the expression data were log2(TPM+1) transformed and the log2FC was defined as median(tumor) – median(normal). p-values were calculated with a one-way ANOVA comparing tumor with normal tissue.

The online tool KMplot (Nagy et al. 2018) was used to analyze different types of survivals and generate Kaplan-meier curves based on gene expression data from microarrays obtained from GEO, caBIG and TCGA. Using the KM plotter online tool, patients were split using the option ‘Auto select best cutoff’ in high or low USP28 gene expression groups. P values for log rank tests of the Kaplan–Meier curves were calculated with the online tool KM plotter. Depmap (version 2020) was used to analyze and visualize Pearson correlation between the genetic expression of USP28 and BRAF/AKT2 in cancer cell lines. p-value and linear regression was calculated by the online tool depmap. https://depmap.org/

Survival of patients with KRAS, EGFR, PIK3CA and BRAF mutated tumors was analysed using the online tool UCSC Xena (Goldman et al. 2020). USP28 gene expression of lung cancer samples (including KRAS, EGFR, PIK3CA and BRAF mutated samples) were obtained from TCGA dataset. Gene expression was downloaded as log2 (norm_count+1). Samples were divided in two groups (High/Low USP28) based on USP28 expression. Expression of USP28 was defined as high when the respective expression levels were higher than the median expression levels of the analysed dataset. Expression of USP28 was defined as low when the respective expression levels were lower than the median expression levels of the analysed dataset. Box plots using TCGA and GTEx data were generated using the online tool BoxPlotR (Spitzer M. et al. 2014) In box plots, the centre line reflects the median and the upper and lower box limits indicate the first and third quartiles. Whiskers extend 1.5× the IQR. For BoxplotR, the data previously download from UCSC Xena was used to generate the graphics, p-values were calculated using two-tailed t-test. Softwares used for this publication are listed in the supplementary table called: Consumables and resources.

### Animal Experiments and histology

All *in vivo* experiments were approved by the Regierung Unterfranken and the ethics committee under the license numbers 2532-2-362, 2532-2-367, 2532-2-374 and 2532-2-1003. All animals are housed in standard cages in pathogen-free facilities on a 12h light/dark cycle with *ad libitum* access to food and water. FELASA2014 guidelines were followed for animal maintenance. Veterinarians supervise the welfare of the animals every day. In presence of pain, stress or suffering, mice were immediately euthanized by cervical dislocation upon Isoflurane anesthesia. The mouse strains used for this publication are listed in the supplementary table called: Consumables and resources.

Adult mice were anesthetized with Isoflurane and intratracheally intubated with 50 μl AAV virus (3 × 10^7^ PFU) as previoulsy decribed (17). Animals were sacrificed by cervical dislocation and lungs were fixed using 5% NBF. For IHC and H×E, slides were de-paraffinized and rehydrated following the previously reported protocol (17). Briefly, IHC slides were subjected to epitope retrieval and blocked in 3% BSA at RT for 1h. Antibody manufacturer instructions were followed for every antibody. But in general, primary antibodies (diluted in 1% BSA) were incubated ON at 4°C followed by three washes with PBS and the subsequent incubation with the DAB secondary antibody for 1 hour at RT. Then, slides were washed twice with 1xPBS for 5 min and stained with the DAB staining solution in 1xPBS. Upon DAB staining, slides were counteracted with hematoxylin and washed three times with 1x PBS for 5 min. Slides were mounted with 200 μl of Mowiol® 40-88 covered up by a glass coverslip. IHC slides were recorded using Pannoramic DESK scanner or using FSX100 microscopy system (Olympus) and analysed using Case Viewer software (3DHISTECH), QuPath software and ImageJ. IF samples were recorded using FSX100 microscopy system (Olympus). Antibodies used in this publication are listed in the supplementary table called: Consumables and resources.

### Organotypic lung tumor slice cultures ex vivo

Lung tumors developed upon endotracheal transplantation of KPL cells as previously described (17) or WT lung tissue from WT C57BL6/J-Rosa26 Sor-CAGG-Cas9-IRES-eGFP animals were explanted and sectioned in slices using the vibratome. Ex vivo slices were relocated in cell culture dishes and maintained in standard cell culture medium (DMEM, 10% FCS) and conditions (37°C, 5% CO2 and 95% relative humidity).

### Sample preparation for mass spectrometry

The sample preparation was performed as described previously. In brief, lysates were precipitated by methanol/chloroform and proteins resuspended in 8 M Urea/10 mM EPPS pH 8.2. Concentration of proteins was determined by Bradford assay and 100 µg of protein per samples was used for digestion. For digestion, the samples were diluted to 1 M Urea with 10mM EPPS pH 8.2 and incubated overnight with 1:50 LysC (Wako Chemicals) and 1:100 Sequencing grade trypsin (Promega). Digests were acidified using TFA and tryptic peptides were purified by tC18 SepPak (50 mg, Waters). 10 µg peptides per sample were TMT labelled and the mixing was normalized after a single injection measurement by LC-MS/MS to equimolar ratios for each channel. A bridge channel was prepared by pooling 3 μg from all 24 samples which were TMT-labeled together and split into two 10 μg samples for each plex, 130 µg of pooled peptides were dried for High pH Reversed-phase fractionation.

### High pH reversed-phase fractionation

Labeled peptide samples were pooled, fractionated into 8 fractions using the High pH Reversed-Phase Peptide Fractionation Kit (ThermoFisher Scientific 84868) according to the manufacturer protocol and dried. Additionally, for label free single shots, 10 µg of peptide is cleaned up with Empore C18 stage tipping and dried right away for shooting.

### LC-MS^3^ proteomics

All mass spectrometry data was acquired in centroid mode on an Orbitrap Fusion Lumos mass spectrometer hyphenated to an easy-nLC 1200 nano HPLC system using a nanoFlex ion source (ThermoFisher Scientific) applying a spray voltage of 2.6 kV with the transfer tube heated to 300°C and a funnel RF of 30%. Internal mass calibration was enabled (lock mass 445.12003 m/z). Peptides were separated on a self-made, 32 cm long, 75µm ID fused-silica column, packed in house with 1.9 µm C18 particles (ReproSil-Pur, Dr. Maisch) and heated to 50°C using an integrated column oven (Sonation). HPLC solvents consisted of 0.1% Formic acid in water (Buffer A) and 0.1% Formic acid, 80% acetonitrile in water (Buffer B).

For total proteome analysis, a synchronous precursor selection (SPS) multi-notch MS3 method was used in order to minimize ratio compression as previously described (McAlister et al., 2014). Individual peptide fractions were eluted by a non-linear gradient from 3 to 60% B over 150 minutes followed by a step-wise increase to 95% B in 6 minutes which was held for another 9 minutes. Full scan MS spectra (350-1400 m/z) were acquired with a resolution of 120,000 at m/z 200, maximum injection time of 100 ms and AGC target value of 4 × 10^5^. The most intense precursors with a charge state between 2 and 6 per full scan were selected for fragmentation within 3 s cycle time and isolated with a quadrupole isolation window of 0.7 Th. MS2 scans were performed in the Ion trap (Turbo) using a maximum injection time of 50ms, AGC target value of 15 × 10^4^ and fragmented using CID with a normalized collision energy (NCE) of 35%. SPS-MS3 scans for quantification were performed on the 10 most intense MS2 fragment ions with an isolation window of 1.2 Th (MS) and 2 m/z (MS2). Ions were fragmented using HCD with an NCE of 65% and analyzed in the Orbitrap with a resolution of 50,000 at m/z 200, scan range of 110-500 m/z, AGC target value of 1.5 ×10^5^ and a maximum injection time of 150ms. Repeated sequencing of already acquired precursors was limited by setting a dynamic exclusion of 60 seconds and 7 ppm and advanced peak determination was deactivated.

### Proteomics analysis

Proteomics raw files were processed using proteome discoverer 2.2 (ThermoFisher). Spectra were recalibrated using the Homo sapiens SwissProt database (2020-03-12) and TMTpro (+304.207 Da) as static modification at N-terminus and Lysines, together with Carbamidomethyl at cysteine residues. Spectra were searched against human database and common contaminants using Sequest HT with oxidation (M) as dynamic modification together with methionine-loss + acetylation and acetylation at the protein terminus. TMTpro (N-term, K) and carbamidomethyl were set as fixed modifications. Quantifications of spectra were rejected if average S/N values were below 5 across all channels and/or isolation interference exceeded 50%. Protein abundances were calculated by summing all peptide quantifications for each protein. Two mixing two plexes, a bridge channel was used additionally. Internal reference scaling (IRS) normalization was performed to obtain proteomics data set across two plexes.

Reactome analysis was performed with PANTHER using the “ Statistical overrepresentation test” tool with default settings. For Reactome analysis and Violin plots, the common proteins significantly dysregulated in EGFR L858R, BRAF V600E and PIK3CA H1047R BEAS-2B respect to DIF. BEAS-2B cells were selected. Proteins were considered significantly downregulated when p-value<0.05. Z-score heatmap visualization was performed using Morpheus (Broad Institute). The maximum and minimal Z-score per row used in heatmaps was calculated using Morpheus (Broad Institute). Volcano plots were generating using the software Instant Clue (Nolte et al. 2018). Venn Diagrams were performed using the online tool: http://bioinformatics.psb.ugent.be/webtools/Venn/. PCA analysis was performed using the online tool CLUSTVIS (Tauno Metsalu and Jaak Vilo 2015). Violin plots were generated using the online tool BoxPlotR (Spitzer M. et al. 2014). For visualization purposes, Excel (Microsoft) and Affinity Designer were used as bioinformatic tools.

## Data and Software availablitiy

Proteomic data is available at PRIDE.

## Contact for reagent and resource sharing

Further information and requests for resources and reagents should be directed to and will be fulfilled by the Lead Contact, Markus E. Diefenbacher.

## Results

### USP28 is expressed in human ‘cell-of-origin’ for NSCLC and upregulated irrespective of lung tumour subtype

Previous work has identified the ‘cell-of-origin’ for NSCLC, in particular adenocarcinoma (BADJ (SPC^+^/CC10^+^) and AT2 (SPC^+^) pulmonary cells)(18, 19), while squamous cell carcinomas, one of the most mutated tumour entities, demonstrated a rather high degree of plasticity and flexibility regarding the required cell of origin(20). Here, the genetic driver combination was the determining factor(21).

We previously observed that genetic loss of USP28 affected tumor burden in a murine *in vivo* NSCLC model(22). To investigate whether this could be attributed to the expression of USP28 in the stem cell/cell of origin compartment of NSCLC, we performed immunohistochemistry against USP28 in human lung tissue samples. High USP28 expression was detectable in the tracheal basal cells (triangular-like shape), BADJ and AT2 cells, the ‘cells-of-origin’ for NSCLC (Figure 1A and 1B). Next, we were intrigued if USP28 expression is altered relative to tumour type and/or grade. To address this question, we analysed publicly available gene expression datasets of NSCLC patients (Figure 1C). *USP28* was found to be upregulated already at an early stage in lung cancer, when compared to wild type tissue (Figure 1C). Furthermore, *USP28* upregulation is a common feature of tumour cells, irrespective of histological or molecular tumour subtype (Figure 1C, S1A and S1B). This was further validated by immunostaining of non-transformed versus tumour samples from NSCLC patients, where a significant increase in USP28 protein abundance was detected already at low grade stages for lung ADC and SCC (Figure 1D and 1E).

**Figure 1:**
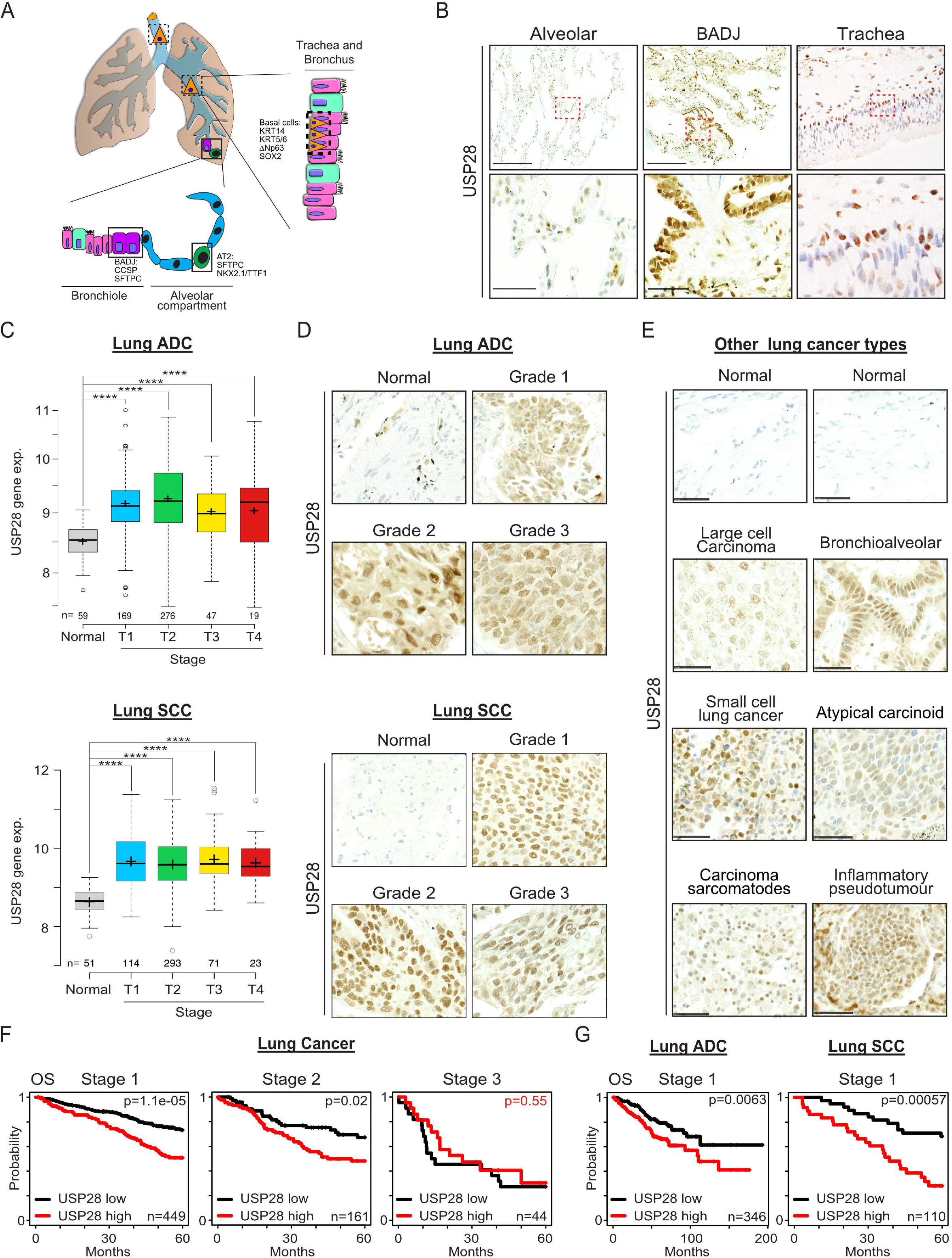
USP28 is expressed in human ‘cell-of-origin’ for NSCLC and upregulated irrespective of lung tumour subtype. A) Schematic representation of the cellular composition of the tracheal, bronchio-alveolar and alveolar compartment. Highlighted are the respective tissue residing stem cells. Trachea = basal cells, Bronchio-alveolar duct junction = BADJ cells, Alveolar compartment = AT2 cells. B) Immuno-histochemical staining of endogenous USP28 in patient lung resected material. Shown are representative images of alveolar, bronchial and tracheal sections. C) Expression of USP28 in non-transformed and NSCLC Adenocarcinoma (ADC) and Squamous Cell Carcinoma (SCC) samples, relative to tumour stage (T1-T4). Public available data. p-values were calculated using two-tailed T-test statistical analysis. Plot was generated using the online tool www.gepia2.cancer-pku.cn. D) Immuno-histochemical staining of USP28 on human NSCLC tissue micro arrays of ADC and SCC origin, ranging from Grade 1 to 3. Where applicable, non-transformed adjacent tissue was included. Shown are representative images per tumour type. E) Immuno-histochemical staining of USP28 on human lung cancer tissue micro arrays of various lung cancer subtypes. Where applicable, non-transformed adjacent tissue was included. lung resected material. Shown are representative images per tumour type. F) Kaplan-Meier plots of NSCLC patient Overall Survival (OS), relative to USP28 expression, at Stage 1 (p=<0.0005), Stage 2 (p=0.02) to Stage 3 (p=0.55). Data was generated using the online tool www.kmplot.com. G) Kaplan-Meier plots of NSCLC ADC and SCC patient Overall Survival (OS), relative to USP28 expression, at Stage 1 (ADC p=0.0063; SCC p=0,00057). Data was generated using the online tool www.kmplot.com. See also Supplementary Figure S1.

Analysing survival data and correlating USP28 expression to tumour stages it became clear that, especially at early stages, USP28^high^ expressing tumours significantly correlated with an overall shortened survival (Figure 1F), and this observation was independent of tumour subtype (Figure 1G).

These data indicated that USP28 is upregulated upon oncogenic transformation and that upregulation appears to be an early event in the tumorigenesis of lung and holds prognostic value.

### USP28 is expressed in murine lung stem cells and required to establish oncogenic transformation in vivo

To investigate that USP28 is required for NSCLC induction and its upregulation occurs at early stages, we utilised CRISPR/Cas9 genetic engineering mouse models of NSCLC(22, 23).

To this end, we analysed the expression pattern of Usp28 by immunohistochemistry in wild-type, non-transformed lungs (Figure 2A). We observed that USP28 showed a comparable expression to human samples (Figure 2A and Figure 1B). Overall Usp28 expression was elevated in putative stem cells, when compared to surrounding/neighbouring differentiated cells (Figure 2A).

**Figure 2:**
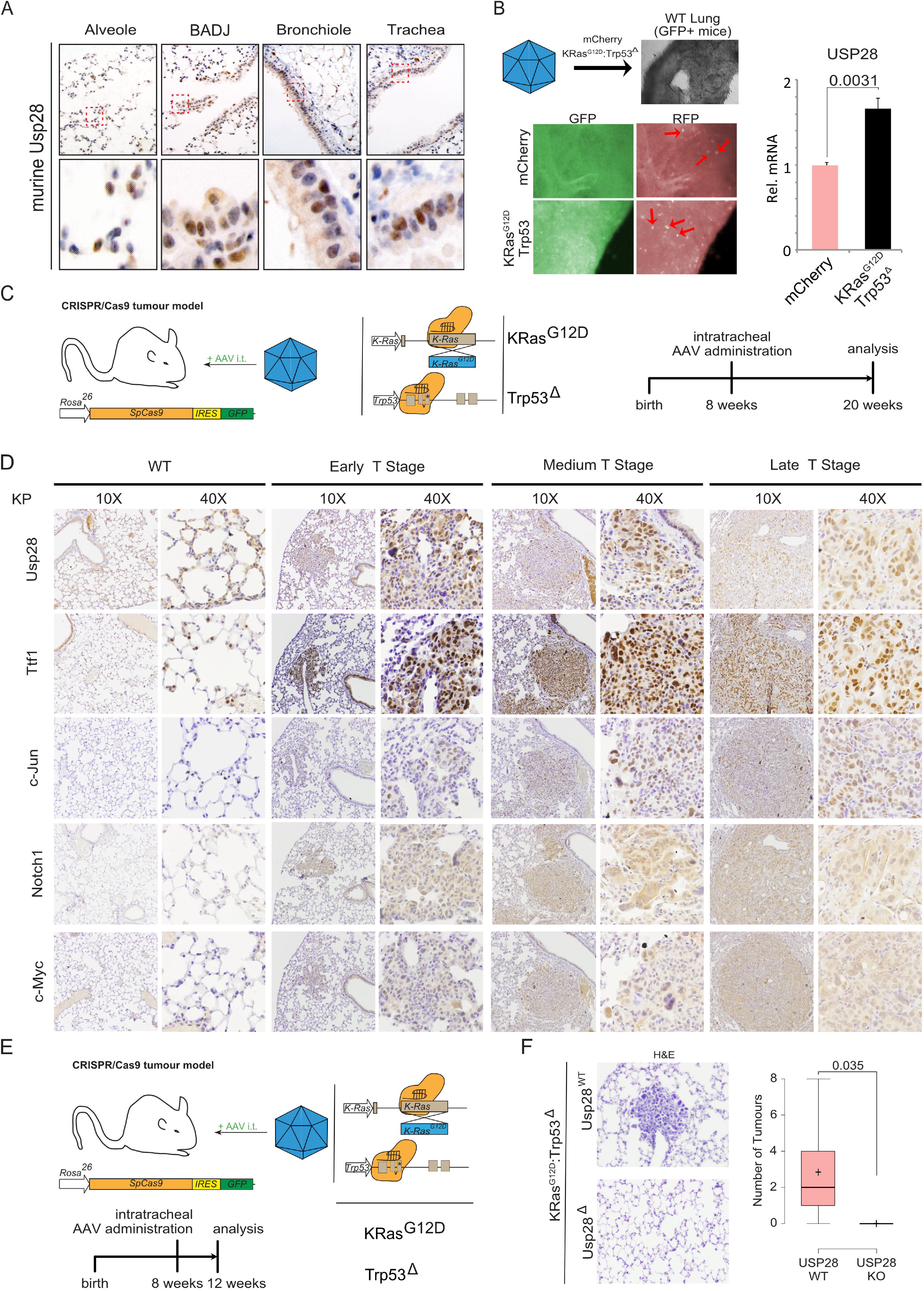
USP28 is expressed in murine lung stem cells and required to establish oncogenic transformation in vivo. A) Immuno-histochemical staining of endogenous Usp28 in murine lung wild type tissue. Shown are representative images of alveolar, bronchial, bronchio-alveolar duct junction and tracheal sections. Red boxes indicate highlighted areas. B) *Ex vivo* onset of oncogenic transduction by CRISPR mediated gene editing and deletion of *Trp53* and mutation of *KRas* to *KRas*^*G12D*^ (KP) upon AAV infection of organotypic lung slice cultures. Lung slice cultures were generated from *C57Bl6/J-Rosa26*^*Sor-CAGG-Cas9-IRES-eGFP*^ mice and AAV encodes mCherry as marker. Fluorescent images of lung slice cultures post infection with AAV. GFP = lung; RFP = AAV infected lung epithelial cells. Tissue sections were harvested and subjected to RNA isolation, followed by RT-PCR analysis of *Usp28* mRNA expression in control and KP infected slices. n= 3 sections each. p-values were calculated using two-tailed T-test statistical analysis. C) Schematic representation of *in vivo* CRISPR gene editing to delete *Trp53* and mutate *KRas* to *KRas*^*G12D*^ (KP) upon intratracheal administering of AAV. Animals are sacrificed 12 weeks post infection. D) Representative images of immuno-histochemical staining against endogenous Usp28, Nkx2-1/Ttf-1, c-Jun, Notch1 and c-Myc in murine lungs infected with either control virus (WT) or KP. Shown are representative tumours spanning grade 1 to 3, with a low and high magnification of individual tumour areas. n=3 E) Schematic representation of *in vivo* CRISPR gene editing to delete *Trp53*, mutate *KRas* to *KRas*^*G12D*^ (KP) or co-delete *Usp28* (KPU) upon intratracheal administering of AAV. Animals are sacrificed 4 weeks post infection. n=6. F) Representative H×E images KP and KPU 4 weeks post infection. Quantification of tumour burden per animal. n=6. p-values were calculated using two-tailed T-test statistical analysis. See also Supplementary Figure S2.

Next, we wondered if the transcriptional upregulation of USP28 already occurs at point of transformation. To address this question, we used an *ex vivo* organotypic lung slice culture model. Here, a non-transformed lung from a C53BL6/J-*Rosa26*^*Sor-CAGG-Cas9-IRES-eGFP*^ mouse(24) was sectioned into 100µm thick sections by using a vibratome and cultured in standard medium. 24 hours post sectioning, slices were infected using an adeno-associated virus (AAV) expressing the fluorescent protein mCherry as infection marker, together with sgRNA to delete *Trp53* and mutate endogenous *Kras* to *Kras*^*G12D*^ (KP, Figure 2B)(22, 23). As a control vector we used an mCherry expressing AAV. 7 days post infection, the lung slices were harvested and mRNA expression of Usp28 analysed using quantitative PCR (Figure 2B). Here, *Usp28* expression was significant increased at an early transformative state, when compared to control virus infected tissue samples.

Next, we analysed the expression of Usp28 and its substrates c-Myc, c-Jun and Notch1 at various grades in murine primary tumours, generated by intratracheal infection with a KP encoding AAV (Figure 2C). Mice were infected at around 8 weeks of age and sacrificed 12 weeks post infection. Tumour grade and type were assessed using histopathological means (H×E, Ttf-1). Already in low grade primary tumours Usp28 and its substrates were significantly upregulated when compared to adjacent, non-transformed lung epithelial tissue (Figure 2D). The increase of Usp28 and the oncogenic transcription factors persisted in higher grade tumours, as seen by immunohistochemistry (Figure 2D), thereby confirming the observation made in patients (Figure 1C, 1D, 1E, S1A and S1B).

Next, we wondered if the upregulation of Usp28 at an early stage is independent of the oncogenic driver (Figure S2A and S2B). For that purpose, we generated murine primary NSCLC using BrafV600E as oncogenic driver. BRAF is genetically altered in 28% of SCC and 25% of ADC lung tumour samples (Figure S2A). To analyse early stage tumours, mice were sacrificed 4 weeks post infection (Figure S2B). Tumour grade and type were determined using H×E and immunohistologic stainings against Ttf1 and Pcna. Already in early stage lung primary tumours, Usp28 was over-expressed when compared to non-transformed tissue (Figure S2C).

Previously we reported that loss of Usp28 affected the induction of lung squamous cancer, and its genetic loss affected overall tumour burden (22). To investigate if loss of Usp28 affects tumour induction at an early stage or just reduces proliferation of transformed cells, next, we infected constitutive Cas9 expressing mice with an AAV virus containing either an sgRNA to delete *Trp53* and mutate endogenous *Kras* to *Kras*^*G12D*^ (KP, Figure 2E) or a virus harbouring additional two sgRNA targeting endogenous Usp28 (KPU, Figure 2E). Mice were sacrificed 4 weeks post infection and while in KP infected animals’ tumour lesions were detectable, in KPU however, no lesions could be observed (Figure 2F).

These data suggest that USP28 is expressed in tumour initiating cells and is required during early stages of lung cancer transformation independent of tumour subtype or oncogenic driver.

### USP28 is increased during early transformation in human BEAS-2B differentiation assay

As it appears that transcriptional upregulation and increased protein abundance of USP28 is a required early event in oncogenic transformation, we used a human cell line system to recapitulate these early events. The immortalised human tracheal cell line BEAS-2B retains the ability to grow as a progenitor like cell, but in a cell-density dependent fashion or under the exposure of fetal calf serum (FCS) can terminally differentiate into a squamous like, pre-oncogenic and highly proliferative cell (Figure 3A)(25, 26). Indeed, culturing these cells under progenitor specific culturing conditions maintained the cells in a ‘spindle-like’ shape as previously reported (Figure 3B; undifferentiated BEAS-2B; BEAS-2B^UD^). Upon FCS exposure, BEAS2B^UD^ cells start to alter their morphology and resemble a squamous phenotype (Figure 3B and 3C; differentiated BEAS-2B; BEAS-2B^DIF^)(25). Furthermore, USP28 protein abundance was increased in BEAS-2B^DIF^ when compared BEAS-2B^UD^ to by immunofluorescence and immunoblotting against endogenous USP28 (Figure 3B and 3C). Not only is USP28 upregulated, but the amount of enzymatically active USP28 was increased as well, indicated by the amount of USP28 bound to a ubiquitin suicide probe/warhead (Figure 3D). Upon pre-oncogenic differentiation, the known USP28 target ΔNp63 increased in protein abundance, as shown by immunoblotting comparing BEAS-2B^DIF^ with BEAS-2B^UD^ (Figure 3B). To investigate if the observed differentiation and the consecutive increase in oncoprotein abundance is USP28 dependent, we exposed BEAS-2B^UD^ to FCS (Figure 3E). Upon serum pulse cells start to differentiate, visualised by brightfield microscopy, and increase the expression of USP28 and its target ΔNp63 (Figure 3E). In order to block USP28, we made use of the published USP28 inhibitor AZ-1(16, 22). Addition of AZ-1 resulted in the degradation of USP28 and reduction in ΔNp63 and cells maintained an undifferentiated morphology (Figure 3E).

**Figure 3:**
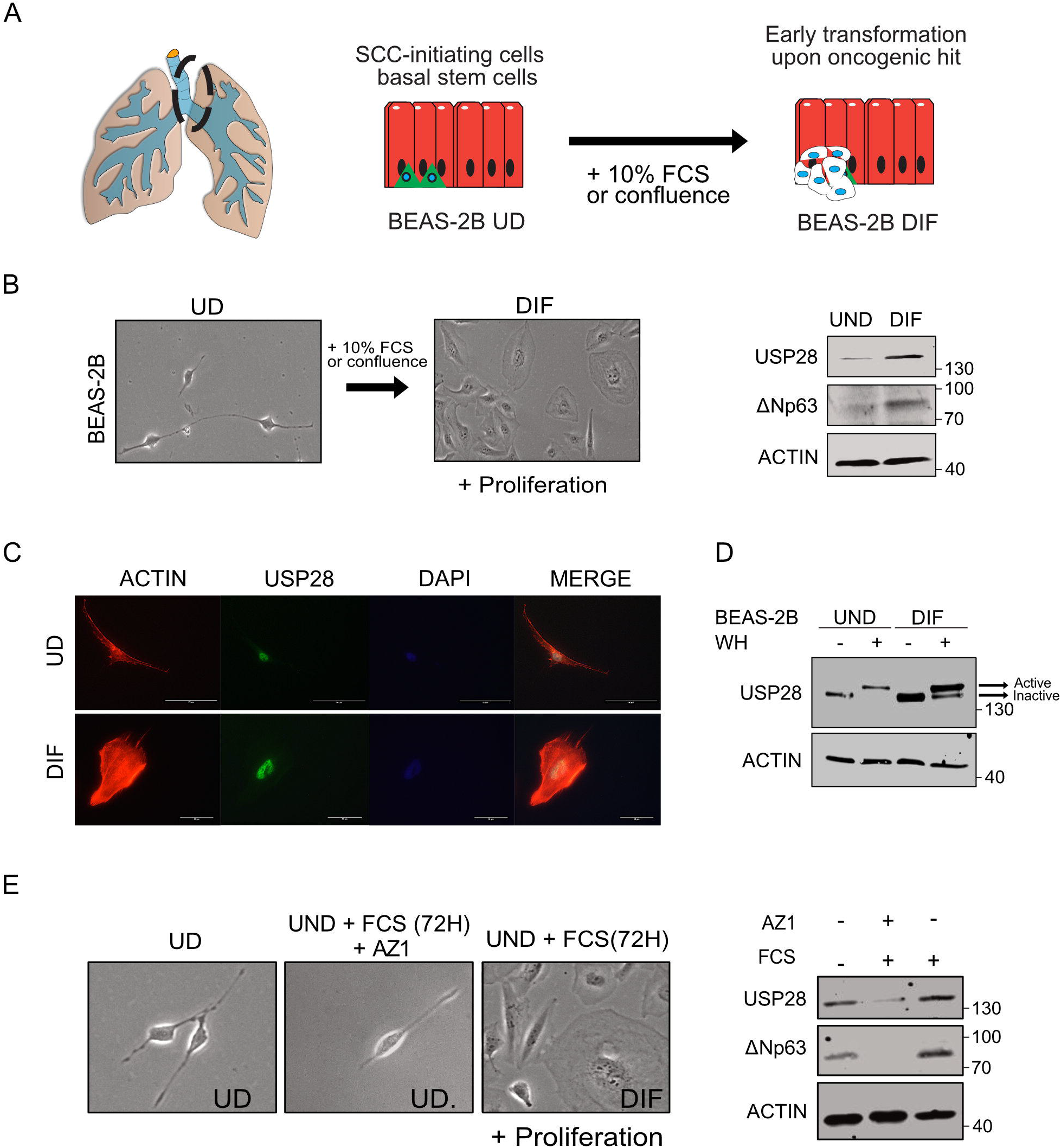
USP28 is increased during early transformation in human BEAS-2B differentiation assay. A) Schematic model of the trans-differentiation by culturing stem cell like, undifferentiated BEAS-2B (BEAS-2B^UND^) in the presence of 10% fetal calf serum to induce squamous like differentiation, BEAS-2B^DIF^. B) Representative brightfield-images of BEAS-2B prior and post serum induced trans-differentiation (72 hours in the presence of 10% FCS, UND to DIF). Immunoblot against endogenous USP28 and ΔNP63 of BEAS-2B^UND^ and BEAS-2B^DIF^. ACTIN served as loading control. Representative blot of n=3. C) Immunofluorescence of endogenous USP28 prior and post differentiation. USP28 in green, ACTIN in red, DAPI as nuclear counterstain. D) Immunoblot against endogenous USP28 in the absence or presence of a Ubiquitin suicide probe (warhead, WH) to assess USP28 enzymatic activity in undifferentiated and differentiated BEAS-2B cells. Upon binding to the activity probe, a shift in molecular weight is observed, indicative of enzymatic activity (see arrows). ACTIN served as loading control. Representative immunoblot of n=3. E) Representative brightfield-images of BEAS-2B prior and post culture in serum induced trans-differentiation conditions, in the presence or absence of the USP28 inhibitor AZ1 (15µM, 72 hours). Immunoblot against endogenous USP28 and ΔNP63 of BEAS-2B^UND^ exposed to either 10% FCS or 10% FCS and 15µM AZ1. ACTIN served as loading control. Representative blot of n=3.

These data demonstrate that USP28 is upregulated during pre-malignant transformation and this leads to an increase of USP28 target oncoproteins.

### Oncogenic transformation of BEAS-2B^DIF^ via EGFR-PI3K-MAPK pathway upregulate USP28 and accelerates tumour cell growth

During the oncogenic transformation process, cells acquire additional alterations, which are the prerequisite to establish a tumour(27). Using public datasets we identified that USP28 expression strongly correlates to the expression of common driver mutations found in NSCLC, encompassing either amplification or mutation of BRAF, EGFR, PI3K and RAS (Figure 4A and 4B, S3A and S3B). HRAS was the exemption in RAS family, as no correlation with USP28 was observed in ADC, however NRAS and KRAS positively correlated with USP28 in ADC human samples (Figure 4B, S3A and S3B). To test this observation, we *in vitro* retroviral transformed BEAS-2B^DIF^ with various oncogenes (BEAS-2B^ONC^; Figure S3C) and investigated the effects on USP28 abundance by immunoblotting and RT-PCR. As a control, we used a virus only encoding Puromycin resistance (Figure S3D and S3E). EGFR WT, EGFRL858R, HRASG12D, BRAFV600E, PIK3CA WT, PIK3CAE545A and PIK3CAH1047R expression was confirmed by immunoblotting and RT-PCR (Figure S3D and S3E). While USP28 was detectable in control transformed cells, the expression of oncogenes further increased USP28 levels and mRNA (Figure 4C and 4D). Not only was the DUB increased, but also the USP28 target protein ΔNp63, along with KRT14, a known ΔNp63 transcriptional target gene (Figure 4C and 4D)(28). It is noteworthy, that the overexpression of wild type PIK3CA/p110 had little to no effect on overall protein increase (Figure 4C and 4D). Here, only the mutant variants E545K and H1047R led to an increase in USP28 and ΔNp63 (Figure 4C and 4D). More studies are required to elucidate the differences between functional mutations and amplification of PIK3CA.

**Figure 4:**
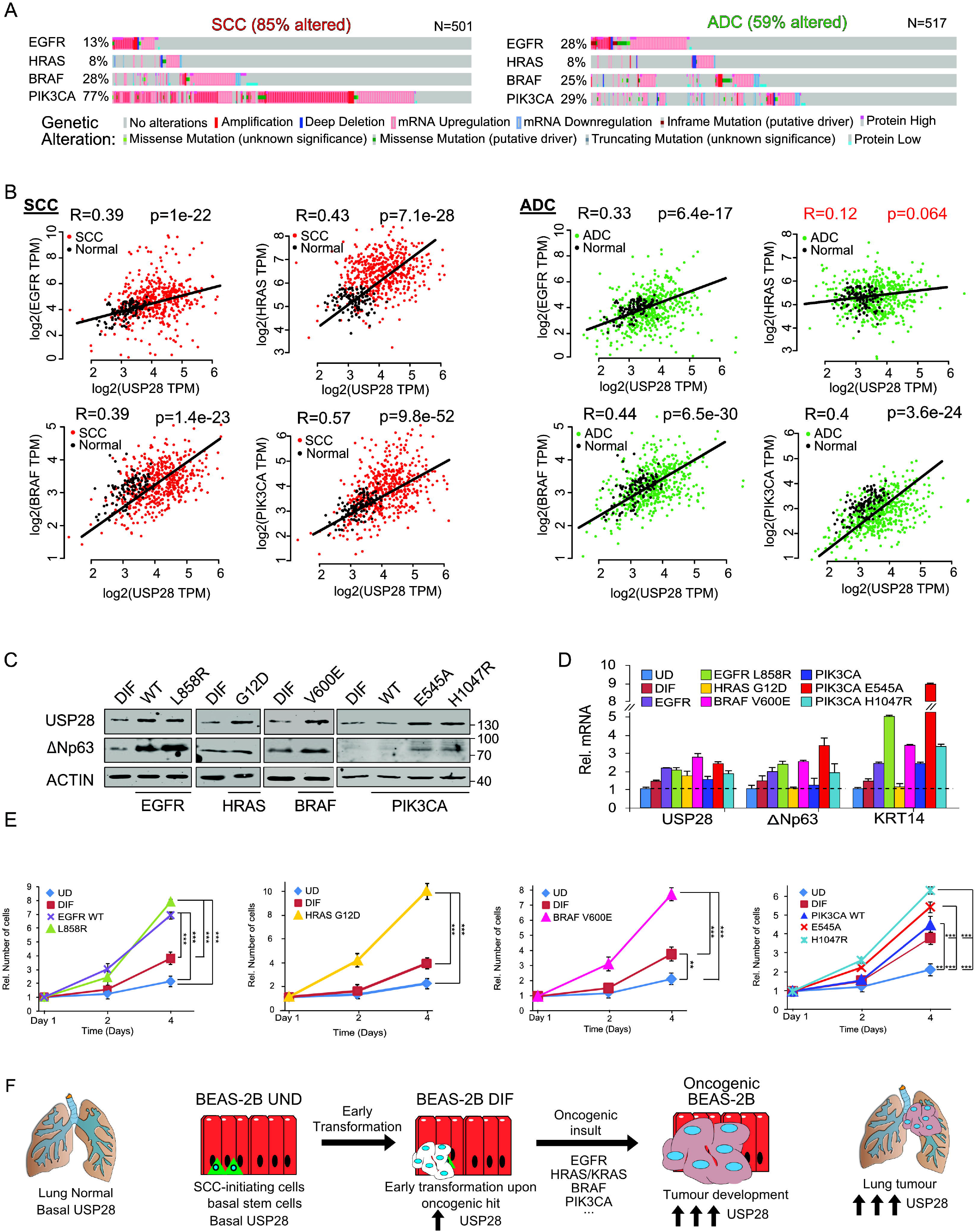
Oncogenic transformation of BEAS-2B^DIF^ via EGFR-PI3K-MAPK pathway upregulate USP28 and accelerates tumour cell growth. A) Frequently occurring genetic alteration in recurring oncogenic drivers found in NSCLC (ADC and SCC). Oncoprints generated with the online tool www.cbioportal.org. B) mRNA expression Spearman’s correlation between USP28 and EGFR, HRAS, BRAF or PIK3CA in NSCLC (SCC and ADC). Correlation and p-value generated with the online tool GEPIA www.gepia2.cancer-pku.cn/. C) Immunoblot against endogenous USP28 and the oncogenic transcription factor ΔNP63 either BEAS-2B^DIF^ or BEAS-2B^DIF^ upon retroviral transduction to express the indicated oncogenes EGFR (wild type (WT) and L858R), HRAS (G12D), BRAF (V600E) and PIK3CA (wild type (WT), E545K and H1047R), respectively. ACTIN served as loading control. Representative immunoblot of n=3. D) RT-PCR of USP28, the SCC transcription factor ΔNP63 and its target Cytokeratin 14 (KRT14) in BEAS-2B^UND^, BEAS-2B^DIF^ or the various BEAS-2B^ONC^ as presented in C). Shown are Log2fold change expression data, relative to ACTIN and normalized to the respective expression in BEAS-2B^UND^ and standard deviation. Shown are mean values and standard deviation of n=3 E) Relative cell numbers and assessment of growth capacity of BEAS-2B^UND^, BEAS-2B^DIF^ or the various BEAS-2B^ONC^ over a total of 4 days. Cell numbers were analyzed every day. Shown are mean values and standard deviation. n=3 experiments. p-values were calculated using two-tailed T-test statistical analysis. F) Schematic model of the various stages of oncogenic transformation, as recapitulated by the trans-differentiation from BEAS-2B^UND^ to BEAS-2B^DIF^, and from BEAS-2B^DIF^ to BEAS-2B^ONC^ (oncogenic transformed BEAS-2B^DIF^). The observed increases recapitulate the increase in USP28 protein abundance as seen in human NSCLC samples. See also Supplementary Figure S3 and S4.

Previous studies reported that USP28 is a direct target of c-JUN and c-MYC (Figure S4A)(29, 30). Since both proto-oncogenes exert an important role during oncogenic transformation and are increased upon EGFR-PI3K-MAPK mediated oncogenic transformation(31), we wondered if the regulation of USP28 in lung cancer depends on these transcription factors as well. Transient transfection of c-MYC and c-JUN increased USP28 protein abundance compared to control plasmid transfection in BEAS-2B cells (Figure S4B). Furthermore, BEAS-2B^DIF^ cells showed higher protein abundance for USP28 and its known substrates c-MYC, c-JUN and NOTCH1 than BEAS-2B^UD^ (Figure S4C) and oncogenic transformation of BEAS-2B^DIF^ further increased USP28, c-MYC, c-JUN and NOTCH1 protein abundance (Figure S4C).

In summary, c-MYC, c-JUN and NOTCH1 are downstream targets of the PI3K-MAPK pathways, and establish a feed forward loop in oncogenic transformed cells, contributing to their increased abundance (Figure S4D). Analysing public datasets revealed that gene expression of the PI3K-MAPK downstream effectors, AKT2 and BRAF, positively correlated with USP28 in lung cancer cell lines and in various tumour entities (Figure S4E). Notably, melanoma and tumours arising in liver, eye and bone showed weak correlation between USP28 and AKT2-BRAF, and for thyroid cancer we observed a negative correlation. This tumour type presents a tumour entity with overall better prognosis and survival(32).

Next, we wondered if oncogenic transformation of BEAS-2B^DIF^ alters cellular growth responses. To investigate potential effects, we compared growth rates of BEAS-2B^UD^, BEAS-2B^DIF^ and BEAS-2B^ONC^ for 4 days (Figure 3E). While the differentiation already enhanced proliferation and is a consequence of enriched proto-oncogene abundance (Figure S4C), upon overexpression of oncogenes, irrespective of oncogenic driver, all generated cell lines demonstrated a significant increase in proliferation, except overexpression of wild type PIK3CA/p110 (Figure 4E).

This observation indicates that USP28 is a downstream target of the PI3K-MAPK pathway (Figure 4F) and is contributing to cellular transformation by stabilising proto-oncogenes (Figure S4D).

### Malignant transformation renders tumour cells dependent on USP28

As USP28 was upregulated in an oncogene dependent fashion, we wondered if the increase in USP28 contributes to the pro-proliferative phenotype. The inducible overexpression of murine USP28 was sufficient to increase the proliferation of BEAS-2B^DIF^ to a comparable extent than PIK3CA mutant cells (Figure 5A). Conversely, using two independent shRNA sequences to target USP28 expression, irrespective of oncogenic driver, loss of USP28 impaired the proliferation of transformed cells (Figure 5B). This was further confirmed using the KRAS G12S mutant lung cancer cell line A549 (Figure S5A). Depletion of USP28 by two independent shRNA sequences strongly reduced the overall abundance of MYC as well the proliferation marker PCNA (Figure S5B). As a consequence, A549 proliferation was significantly reduced (Figure S5C). Next, we wondered if the small molecule inhibitor AZ1, which impairs USP25/28 enzymatic activity, affects proliferation of oncogenic transformed BEAS-2B. Furthermore, we wanted to address if the inhibitor shows selectivity towards specific oncogenic drivers. To this end, BEAS-2B^DIF^ and BEAS-2B^ONC^ cells were grown in the presence of increasing concentration of AZ1, followed by calculating the half maximal inhibitory concentration (IC_50_) by assessing cell numbers. It was revealed that EGFRL858R and BRAFV600E transduced BEAS-2B^ONC^ cells tolerated ∼16µM AZ1, while cells transformed by PIK3CAH1047R required ∼20µM (Figure S5D). Given the rather comparable IC_50_ concentrations, we treated BEAS-2B^DIF^ and BEAS-2B^ONC^ cells with 15 µM AZ1 for 24 hours, followed by immunoblotting against USP28 and its substrates NOTCH1, c-MYC and c-JUN (Figure 5C). Upon exposure to AZ1, all cell lines showed a marked reduction in the protein abundance of USP28 and its target substrates (Figure 5C and 3E). Next, we investigated if AZ1 impairs cell proliferation or increases cell death in a dose dependent fashion, and if AZ1 affects all cells or is limited to oncogenic transformed cells (Figure 5D and 5E). Cells were grown in the presence of increasing concentrations of AZ1 for 72 hours, followed by *in vivo* exposure to propidium iodide (PI) as a marker for dead cells (Figure 5E) and immune-fluorescent staining of the proliferative marker Ki67 (Figure 5D). While the non-oncogenic BEAS-2B^DIF^ showed weak reduction in cell proliferation and mild increase in nuclear PI positive cells, BEAS-2B^ONC^ cells, irrespective of oncogenic driver, significantly reduced cell proliferation in an AZ1 concentration dependent fashion (Figure 5D). Additionally, when BEAS-2B^ONC^ cells were exposed to AZ1 concentrations reaching or exciding the oncogene corresponding IC_50_, PI incorporation was significantly enriched (Figure 5D and 5E). Surprisingly, BRAFV600E transformed BEAS-2B demonstrated a high degree of sensitivity towards AZ1 (Figure 5E), as it was previously reported that, in melanoma, loss of USP28 was required to induce oncogenic transformation and resistance.

**Figure 5:**
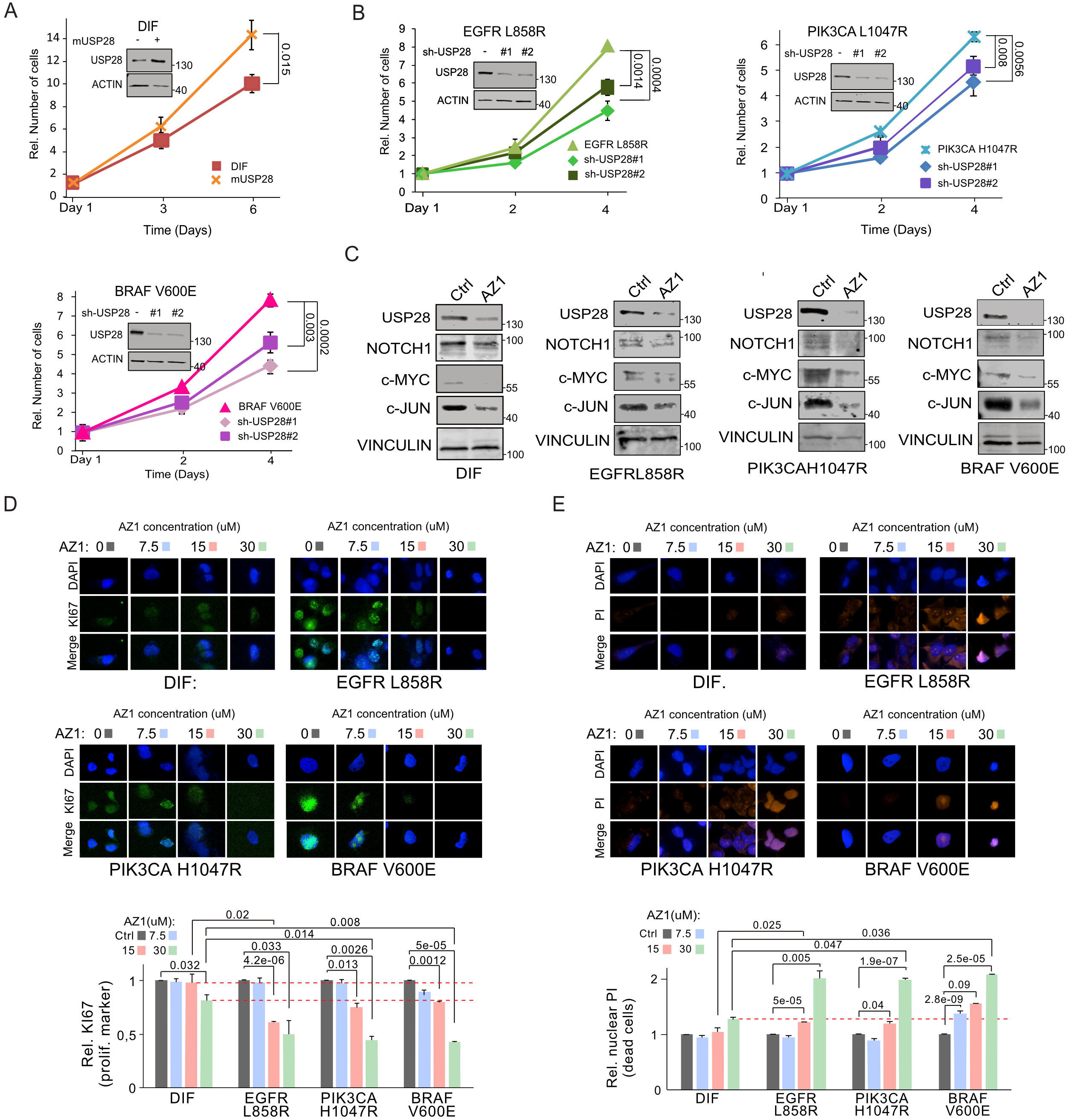
Malignant transformation renders tumour cells dependent on USP28. A) Immunoblot showing protein abundance of USP28 in BEAS-2B^DIF^ upon lentiviral transduction with either a control or a doxycycline inducible overexpression of murine Usp28. BEAS-2B^DIF^ control and mUsp28 cells were cultured in the presence of 1µg/ml doxycycline for 72 hours prior to immunoblotting. ACTIN served as loading control. For growth analysis, cells were pre-cultured for 72 hours in the presence of 1µg/ml doxycycline, followed by re-seeding and counting of cells at day 1, day 3 and day 6. Shown are mean values and standard deviation of n=3. p-values were calculated using two-tailed T-test statistical analysis. B) Immunoblot showing protein abundance of USP28 in oncogenic transduced BEAS-2B^ONC^ (EGFRL858R, PIK3CA L1047R and BRAFV600E) upon lentiviral transduction with either a control or two individual constitutive shRNA targeting USP28. ACTIN served as loading control. For growth analysis, cells were seeded at equal cell density and counted at day 1, day 2 and day 4. Shown are mean values and standard deviation of n=3. C) Immunoblots against endogenous USP28 and its substrates NOTCH1, c-MYC and c-JUN in either BEAS-2B^DIF^ or oncogenic transduced BEAS-2B^ONC^ (EGFRL858R, PIK3CA L1047R and BRAFV600E) upon exposure to either DMSO or 15µM AZ1 for 24 hours. VINCULIN served as loading control. D) Immunofluorescence of Ki-67 expression in BEAS-2B^DIF^ or oncogenic transduced BEAS-2B^ONC^ (EGFRL858R, PIK3CA L1047R and BRAFV600E) cultured in the presence of increasing concentrations of AZ1 (0 (DMSO), 7.5µM, 15 µM, 30 µM) for 72 hours to assess effects in cell proliferation. Shown are representative cells. Quantification of Ki-67 expression in 30-45 20x fields from independent wells per condition. DAPI served as nuclear marker. Shown are mean values and standard deviation. p-values were calculated using two-tailed T-test statistical analysis. E) Immunofluorescence of propidium iodide (PI) incorporation in BEAS-2B^DIF^ or oncogenic transduced BEAS-2B^ONC^ (EGFRL858R, PIK3CA L1047R and BRAFV600E) cultured in the presence of increasing concentrations of AZ1 (0 (DMSO), 7.5µM, 15 µM, 30 µM) for 72 hours to assess effects on cell survival and apoptosis. Shown are representative cells. Quantification of PI positive cells in 30-45 20x fields from independent wells per condition. DAPI served as nuclear marker. Shown are mean values and standard deviation. p-values were calculated using two-tailed T-test statistical analysis. See also Supplementary Figure S5.

Overall, our data demonstrated that USP28 is required to maintain tumour cell proliferation and survival *in cellulo*, hence, tumour cells become addicted to USP28.

### Inhibition of USP28 via AZ1 ‘resets’ the proteome of oncogenic transduced cells towards a ‘non-oncogenic’ state and induces pro-apoptotic signatures

To gather further insights into how targeting of USP28 via AZ1 affects oncogenic transduced BEAS-2B cells, we conducted mass spectrometric analysis and compared the proteome of control and oncogenic transduced cells upon exposure to AZ1 (Figure S6A). Principal component analysis identified that oncogenic transduction resulted in distinct changes of the proteome (Figure 6A and Figure S6B). Not only were the proteomes of BEAS-2B^onc^ different when compared to non-oncogenic cells, but the oncogenic driver used established distinct proteomic patterns (Figure 6A). Upon exposure to AZ1, the proteomes of BEAS-2B^onc^, however, significantly changed and clustered with the expression of non-oncogenic BEAS-2B (Figure 6A and S6C). Analysing the proteome of the three BEAS-2B^onc^ cell lines post AZ1 exposure revealed that a set of proteins dysregulated during oncogenic transformation was commonly affected in an AZ1 dependent fashion; 45 proteins were decreased and 29 proteins commonly increased in BEAS-2B^onc^ (Figure 6B, 6C and S6D). Proteins upregulated in EGFRL858R transduced BEAS-2B were repressed upon exposure to AZ1, while proteins decreased during the course of oncogenic transformation enriched upon blockage of USP28 activity (Figure 6C, 6D and S6D). Addition of 15 µM AZ1 for 72 hours, however, not only reduced the abundance of proto-oncogenes, but re-shaped global protein abundance, closer resembling non-oncogenic BEAS-2B (Figure 6C and 6D). Inhibition of USP28 via AZ1 in oncogenic transduced BEAS-2B significantly repressed the abundance of proteins involved in negative regulation of the ubiquitin proteasome system, decreased RTK/growth factor signalling and vesicle transport, while oncogenic BEAS-2B upregulated proteins involved in differentiation, immune signalling, apoptosis and necrosis (Figure 6E).

**Figure 6:**
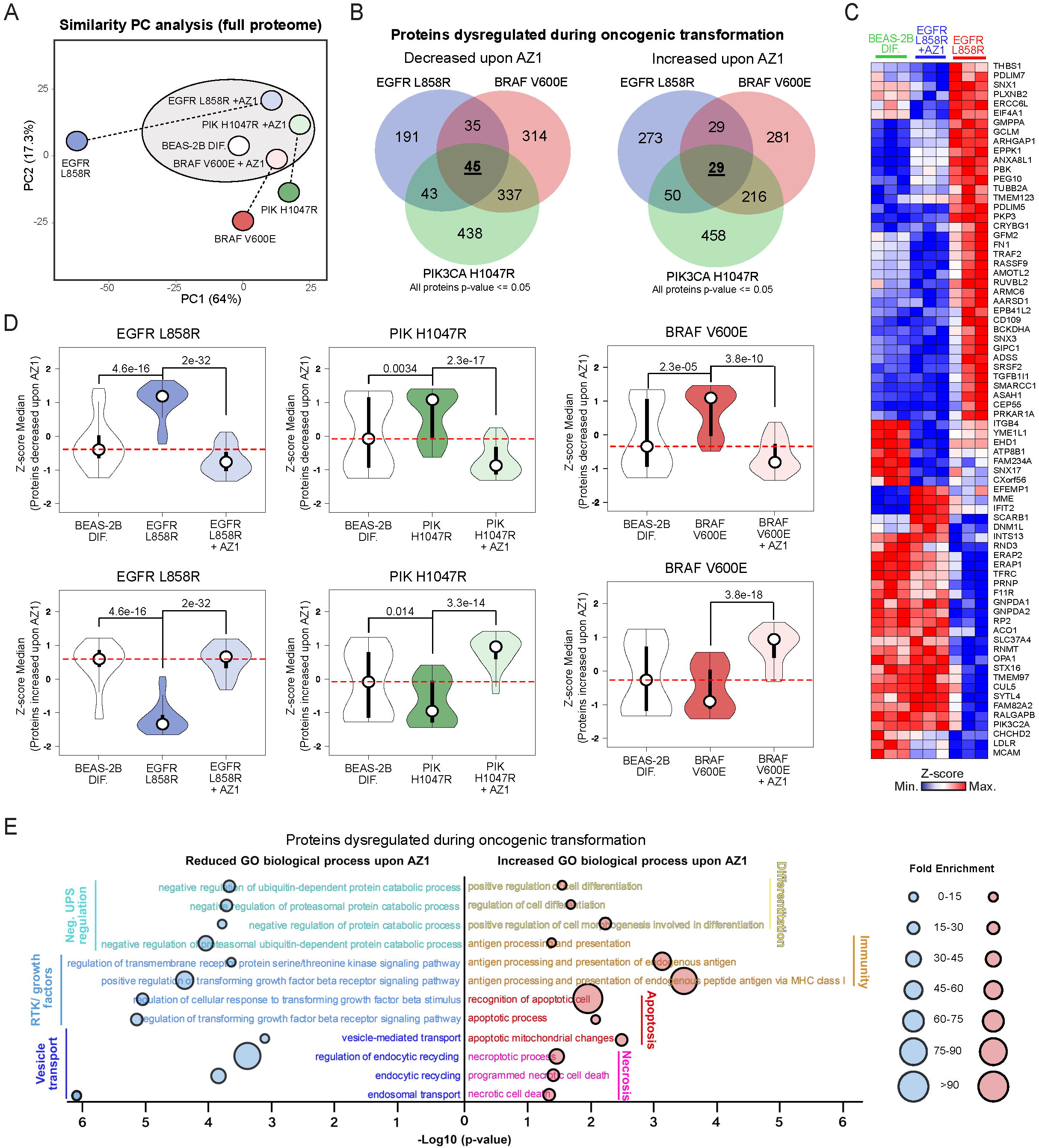
Inhibition of USP28 via AZ1 ‘resets’ the proteome of oncogenic transduced cells towards a ‘non-oncogenic’ state and induces pro-apoptotic signatures. A) Principal Component Analysis (PCA) of the whole proteome of BEAS-2B^DIF^ and BEAS-2B^ONC^ (EGFRL858R, PIK3CA L1047R), treated with either 15 µM AZ1 for 72 hours or exposed to control solvent (DMSO). N=3 samples. Analysis generated with Clustvis online tool (https://biit.cs.ut.ee/clustvis). B) Venn diagram illustrating proteins changed upon exposure of BEAS-2B^ONC^ (EGFRL858R, PIK3CA L1047R) to 15 µM AZ1 for 72 hours. Highlighted are numbers of proteins dysregulated during oncogenic transformation (BEAS-2B BEAS-2B^ONC^) and decreased or increased upon AZ1 exposure. Discreet or common deregulated protein numbers are indicated within the corresponding overlapping graphs. Common, oncogenic driver independent number of deregulated proteins are highlighted in the center. Analysis of n=3 samples per oncogenic driver. C) Heatmap of proteins identified in B) for BEAS-2B^DIF^, BEAS-2B^EGFRL858R^ and BEAS-2B^EGFRL858R^ treated with 15 µM AZ1 for 72 hours. Shown are n=3 experiments and data presented as Z score values. Red= high Z-score protein abundance, blue = low Z-score protein abundance. D) Violin plots illustrating changes of BEAS-2B^DIF^ and BEAS-2B^ONC^ pre- and post-treatment with 15 µM AZ1 for 72 hours for decreased or increased proteins identified in B). p-values were calculated using two-tailed T-test statistical analysis.For Violin plot, white circles show the medians; box limits indicate the 25th and 75th percentiles as determined by R software; whiskers extend 1.5 times the interquartile range from the 25th and 75th percentiles; polygons represent density estimates of data and extend to extreme values. Violin plot were generated with BoxplotR. http://shiny.chemgrid.org/boxplotr/ E) GO biological process significantly reduced or increased in BEAS-2B^ONC^ upon 15 µM AZ1 for 72 hours. The analysis was performed with the proteins identified in B). The analysis was performed using the online tool Panther. http://www.pantherdb.org See also Supplementary Figure S6.

Hence, acute inhibition of USP28 via AZ1 decreases the amount of proto-oncogenes, thereby affecting global protein abundance. USP28 is required to accommodate oncogenic transformation and to suppress anti-proliferative and pro-apoptotic signatures.

### USP28 inhibition potentiates targeted molecular therapy

To investigate if USP28 is a putative prognostic marker for NSCLC, we analysed public available datasets and observed that USP28 expression significantly correlates with progression free survival (PFS, p=4.6e-05), Figure 7A and S7A). In the Post Progression Survival cohort, elevated expression of USP28 directly correlated with shortened survival (overall and Stage I, S7A and S7B). These data suggest that USP28 levels strongly determine the survival of the NSCLC patients in response to therapy.

**Figure 7:**
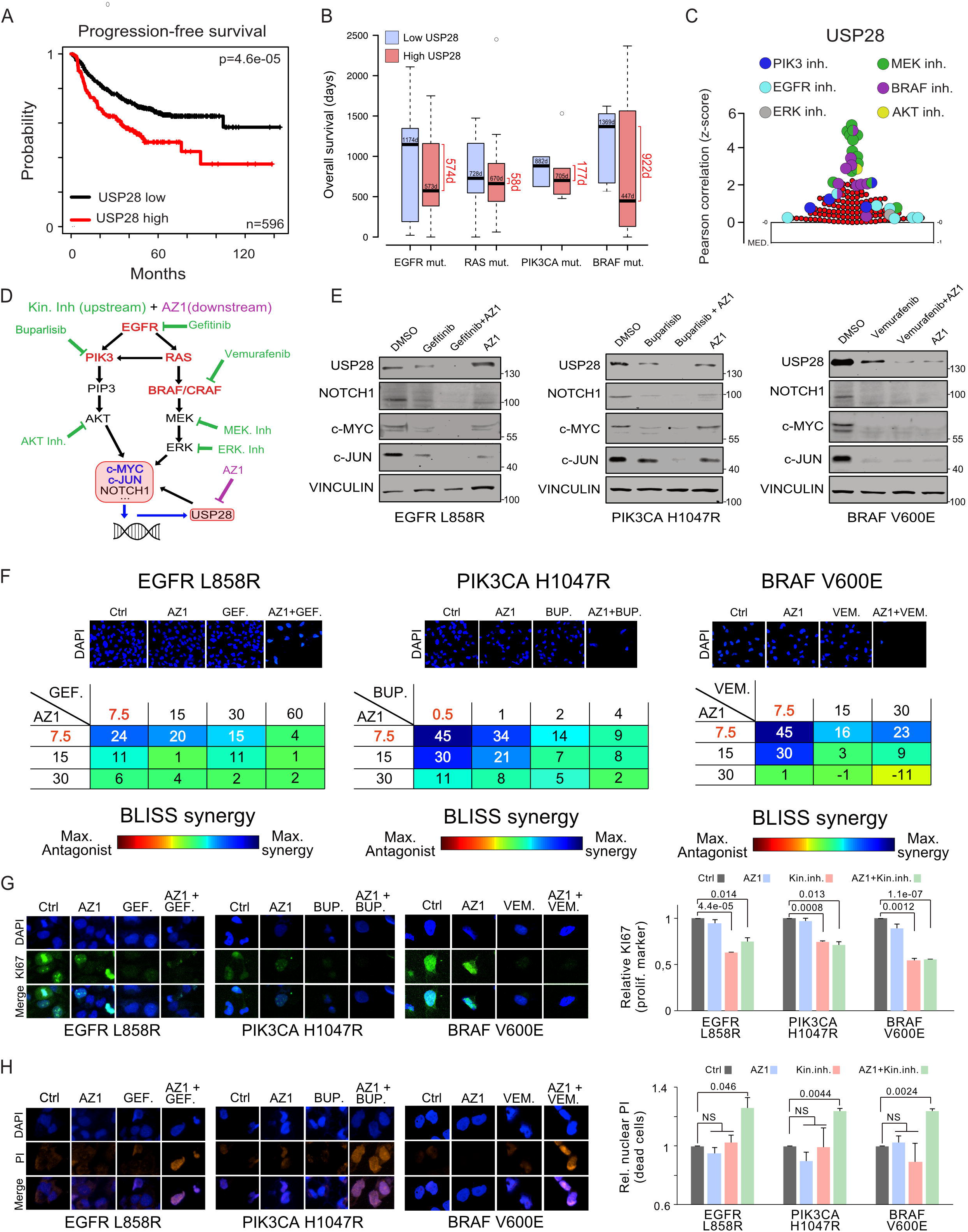
USP28 inhibition potentiates targeted molecular therapy. A) Kaplan Meier plot of Progression free survival (PFS) of NSCLC patients relative to USP28 mRNA expression. n=596 samples, p=<0.0005. Data generated with the online tool www.kmplot.com. B) Analysis of public available datasets analysing USP28 mRNA expression and Overall Survival (OS) in patients with mutations in the oncogenic driver EGFR, RAS, PIK3CA or BRAF. Samples were divided in two groups based on USP28 mRNA expression: High USP28 (higher than the median USP28 expression) and Low USP28 (lower than the median USP28 expression). Survival days was determined for both groups. Data was obtained from the online tool https://xena.ucsc.edu/. In box plots, the centre line reflects the median and the upper and lower box limits indicate the first and third quartiles. Whiskers extend 1.5× the IQR. C) Pearson correlation between sensitivity of EGFR-PI3K-MAPK inhibitors and USP28 expression in human NSCLC cell lines. Data was generated with the online tool Cancer Therapeutics Response Portal V2 (https://portals.broadinstitute.org/ctrp/). In plos, the lower line reflects the median and the upper box limit indicates the first quartile. Whiskers extend 1.5× the IQR. D) Schematic representation of EGFR-PI3K-MAPK pathway analysed in this study and potential pathway interference opportunities by EGFR-PI3K-MAPK and AZ1 inhibitors. Green = EGFR-PI3K-MAPK inhibitors. Violet = AZ1. E) Immunoblots of BEAS-2B^ONC^ (EGFRL858R, PIK3CA L1047R and BRAFV600E) cultured in the presence of either control solvent (DMSO), pathway specific inhibitors (EGFR: 20 µM Gefitinib; PIK3CA: 1µM Buparsilib; BRAF: 20 µM Vemurafenib), 15 µM AZ1 or a combination thereof for 24 hours. F) BLISS synergism score of BEAS-2B^ONC^ (EGFRL858R, PIK3CA L1047R and BRAFV600E) cultured in the presence of either control solvent (DMSO), pathway specific inhibitors (EGFR: Gefitinib; PIK3CA: Buparsilib; BRAF: Vemurafenib), AZ1 and combination thereof for 72 hours at indicated concentrations. Shown are representative DAPI images of cells 72 hours post culture in the presence of DMSO, single treatment with 7.5 µM AZ1, 7.5 µM Gefitinib, 0.5µM Buparlisib, 7.5µM Vemurafenib or combination of AZ1 with the respective personalized molecular therapy for 72 hours. Synergism was calculated by cell quantification of 30-45 20x fields from independent wells in control (DMSO) and the indicated treated conditions. G) Immunofluorescence of Ki-67 expression in oncogenic transduced BEAS-2B^ONC^ (EGFRL858R, PIK3CA L1047R and BRAFV600E), cultured in the presence of DMSO, single treatment with 7.5 µM AZ1, 7.5 µM Gefitinib, 0.5µM Buparlisib, 7.5µM Vemurafenib or combination of AZ1 with the respective pathway inhibitor for 72 hours, to assess effects in cell proliferation. Shown are representative cells. Quantification of Ki-67 expression in 30 to 45 20x fields from different wells in control (DMSO) and treated conditions. DAPI served as nuclear marker. Shown are mean values and standard deviation. p-values were calculated using two-tailed T-test statistical analysis. H) Immunofluorescence of propidium iodide (PI) in vivo incorporation of oncogenic transduced BEAS-2B^ONC^ (EGFRL858R, PIK3CA L1047R and BRAFV600E), cultured in the presence of DMSO, single treatment with 7.5 µM AZ1, 7.5 µM Gefitinib, 0.5µM Buparlisib, 7.5µM Vemurafenib or combination of AZ1 with the respective pathway inhibitor for 72 hours, to assess effects in cell proliferation. Shown are representative cells. Quantification of PI positive cells in 30 to 45 20x fields from different wells in control (DMSO) and treated conditions. DAPI served as nuclear marker. Shown are mean values and standard deviation. p-values were calculated using two-tailed T-test statistical analysis. See also Supplementary Figure S7.

Since NSCLC is a genetically very heterogeneous tumour type, next, we wondered if patient survival depended on the combination of oncogenic driver and the expression of USP28 (Figure 7B and S7C). Indeed, the analysis of public available data leads to the suggestion that, irrespective of oncogenic driver, an increased expression of USP28 significantly shortens survival for tumours driven by mutations in the oncogenes EGFR (Δ574 days), PIK3CA (Δ177 days) or BRAF (Δ922 days), while in tumours driven by mutations in genes of the RAS family, USP28 had very little effect on patient survival (Δ58 days, Figure 7B). These data suggest that USP28 is a suitable prognostic marker and oncogene in NSCLC.

As cancer panel sequencing is implemented in the clinics and pathway-specific inhibitors available, we were wondering if disruption of the oncogenic pathways would directly affect USP28 and hence, the abundance of its downstream effectors. Analysis of publicly available data regarding putative drug sensitivity of tumour cells in direct correlation to USP28 expression scored the PIK3-, EGFR- and MAPK pathway as top hits (Figure 7C), directly confirmed our experimental data regarding USP28 expression relative to oncogenic drivers. Since several potent pathway inhibitors are readily available (Figure 7D), we wondered if targeted therapy would synergize with targeted inhibition of USP28 via AZ1. To this end, we exposed our BEAS-2B^ONC^ cell lines to either a selective inhibitor (EGFRL858R= Gefitinib; PIK3CAH1047R= Buparlisib; BRAFV600E= Vemurafenib), AZ1, or a combination thereof for 24 hours, followed by immunoblotting against USP28 and its substrates (Figure 7E). Selective pathway interference was evaluated by immunoblotting (Figure S7D). Monotherapy via selective pathway inhibitors affected downstream signalling cascades, reduced the abundance of USP28, led to a reduction in the protein levels of NOTCH1, c-MYC and c-JUN and reduced cell viability (Figure 7E and S7E). Similar effects were observed by administering AZ1; treated cells showed a reduction in USP28 abundance, along with reduced protein levels of its substrates and decreased viability (Figure 7E and S7E). Combinatorial treatment, however, significantly reduced the amount of USP28 and diminished the abundance of NOTCH1, cMYC and cJUN (Figure 7E). Not only does co-treatment with AZ1 sensitize BEAS-2B^ONC^ cells to targeted therapy, but it synergises with Gefitinib, Bupalislib and Vemurafenib, as seen by viability assays in BEAS-2B^ONC^ cells (Figure 7F). The synergistic effect of AZ1 and targeted therapy does not only stem from impairment of tumour cell proliferation, but also leads to onset of cell death, as seen by Ki67 expression and PI incorporation experiments of BEAS-2B^ONC^ cells cultivated for 72 hours in the presence of the single compounds or combinations thereof at synergistic concentrations (AZ1: 7.5 µM, Gefitinib: 7.5 µM; Buparlisib: 0.5 µM; Vemurafenib: 7.5 µM; Figure 7G, H and S7E). Here, exposure to 7.5 µM AZ1 did not affect proliferation nor induced cell death (Figure 7G and H). While monotherapy with targeted inhibitors led to a reduction in proliferation, it did not result in cell death, as seen by loss of Ki67 expression and the absence of PI incorporation (Figure 7G and H). The combinatorial treatment, however, resulted in a significant increase in PI positive cells, while maintaining a proliferative block.

USP28 therefore, functions as a signal amplifier downstream of various oncogenic signalling cascades and is required to allow oncogenic transformation. Its inhibition potentiates targeted therapy by impairing the protein stability of oncogenic transcription factors, required to maintain tumour cell proliferation, and presents a promising target for drug development.

## Discussion

Oncogenic transformation of somatic cells is a multistage process frequently starting with the inactivation of tumor suppressors and subsequent gain of activating mutations in oncogenic drivers, such as members of the PI3K or MAPK family. These changes result in the increased abundance of proto-oncogenes, such as c-MYC^(33)^, JUN^(34)^ or NOTCH, driving cell proliferation, dedifferentiation, metabolic changes, DNA damage control, immune evasion and proteostatic stress management, the ‘hallmarks of cancer’(23).

Cells undergoing transformation partially counter this intrinsic stress by re-adjusting the UPS. USP28 is involved in the control of a plethora of biological processes. It is involved in the regulation of cell proliferation and differentiation via its ability to regulate the abundance of the proto-oncogenes, such as c-MYC, c-JUN or NOTCH (Figure 8A) (29). It is involved in transcriptional control via regulating the abundance of the histone modifier LSD1/KDM1A(35) and it is part of the DNA damage machinery, where it interacts with ATM(36), CLSPN(37), or TP53BP1(38).

**Figure 8:**
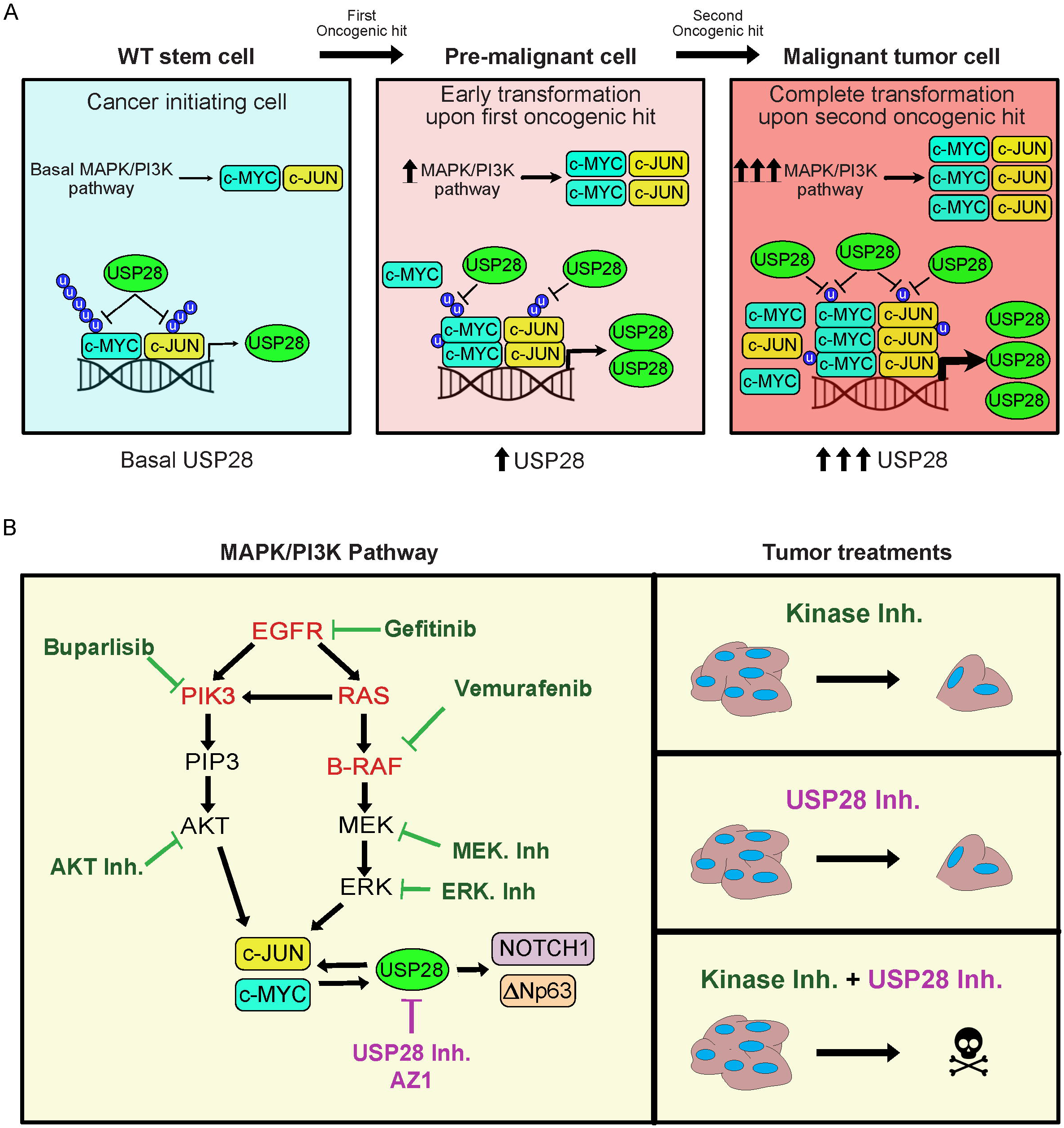
USP28 enables oncogenic reprograming of respiratory cells during early transformation and its inhibition potentiates targeted molecular therapy. A) Schematic representation of the mechanism presented in this manuscript. Oncogenic transformation via EGFR-PI3K-MAPK pathway increases USP28 transcription via c-MYC/ c-JUN. USP28 stabilizes the oncoproteins c-MYC/ c-JUN stablishing a direct feedback loop. B) Schematic representation of the synergy between EGFR-PI3K-MAPK targeted molecular therapies and AZ1 presented in this manuscript.

Overall, USP28 functions as a proto-oncogene and contributes to establishing the hallmarks of cancer in cells undergoing oncogenic transformation, at least in lung(22). Here, USP28 is expressed in the stem cell niche and elevated abundance detected in ‘cells or origin’. Its upregulation is an early event, as already low grade tumours showed enhanced abundance for this particular DUB, and its expression coincided with shortened overall survival. In a cellular multi-stage transformation model, we were able to recapitulate the stepwise increase in USP28 abundance and enhanced activity, which occurred independent to the oncogenic driver present. USP28 is a transcriptional target of its own substrates(29), and via a feed forward loop, increased in cancer compared to normal. Irrespective of oncogenic driver, interference with USP28 abundance or activity suppressed tumour cell growth and survival of transformed lung cells *in vitro* and *in vivo*. Inhibition of USP28 via the small molecule inhibitor AZ1 restored the proteome of oncogenic transformed BEAS-2B cells towards a pre-malignant state, demonstrating that interference with USP28 abundance and activity has a far-reaching biological impact. USP28 is a downstream target of RTK signaling cascades and required to establish oncogenic transformation by its ability to control the abundance of proto-oncogenes. The inhibition of the DUB not only repressed tumour cell growth but led to a pro-apoptotic phenotype, predominantly in oncogenic transformed cells, while control cells were not affected. Here, the cell cycle was affected, leading to a reduction in proliferation.

This is in stark contrast to two recent reports. In a small cohort of melanoma patients, where tumours are predominantly driven by mutant BRAF(V600E), USP28 is genetically lost(39). These USP28 mutant patients present enhanced MAPK signaling via hyperstabilisation of RAF family members and resistance to BRAF inhibitors. Here, loss of USP28 presents a negative survival marker. The second study identified that, in melanoma cells, USP28 is cleaved by caspase 8 to overcome G2/M cell cycle arrest in a TP53 dependent fashion(40). Loss of USP28 in tumour cells is favored as it results in TP53 protein destabilization, thereby establishing a switch of cell fate, from apoptosis towards mitosis. Hence, USP28 functions as a tumour suppressor in melanoma(41). In line with these reports, we could not detect a correlation between USP28 and BRAF expression in human skin cancer samples. This could be indicating that this tumour entity indeed does not rely on the DUB and hence, alternative mechanisms deviating from our observation in lung and SCC, are possible.

In lung, and as previously reported for squamous cell carcinoma(22), targeting USP28 presents a suitable lever for therapeutic engagement, as one could propose that at least in lung, tumour cells become addicted to USP28. In line with this hypothesis, we indeed did observe that loss of USP28 reduced oncogenic cell proliferation and its genetic loss impaired tumour onset *in vivo*. The inhibition of USP28, via a small molecule inhibitor AZ1, induced pro-apoptotic signaling in cells expressing potent oncogenic driver mutants. In lung, the expression of USP28 directly correlated with shortened patient survival, irrespective of oncogenic driver. As USP28 functions as an ‘amplifier’ downstream of RTKs, its inhibition cooperated with personalized targeted therapy against specific oncogenic driver mutations (Figure 8B).

Overall, our data suggest that targeting USP28 protein abundance and activity already at an early stage, therefore, is a promising strategy for the treatment of lung tumours in combination to personalized targeted therapy.

## Supporting information

Consumables and resources

## Acknowledgements

We are grateful to the animal facility and Barbara Bauer at the Biocenter, University Würzburg. C.P.G. and O.H. are supported by the German Cancer Aid via grant 70112491, M.R. is funded by the DFG-GRK 2243 and IZKF B335. M.E.D. and M.R. are funded by the German Israeli Foundation grant 1431. T. F. is funded by the IZKF program Z2/CS-1.

