## Supplementary material for "USP28 enables oncogenic transformation of respiratory cells and its inhibition potentiates molecular therapy targeting mutant EGFR, BRAF and PI3K": Consumables and resources

| **1^st^ Antibodies** | **Company** | **Identifier** |
| --- | --- | --- |
| USP28 | Sigma-Aldrich | HPA006778 |
| B-ACTIN (C4) | Santa Cruz | sc-47778 |
| VINCULIN | Sigma-Aldrich | V9131 |
| P63 | Biolegend | 619001 |
| P63 | Bimake | A5182 |
| c-MYC (N-262) | Santa Cruz | sc-764 |
| c-JUN (G-4) | Santa Cruz | sc-74543 |
| NOTCH1 | Bimake | A5176 |
| EGFR (A10) | Santa Cruz | sc-373746 |
| HRAS (259) | Santa Cruz | sc-35 |
| BRAF (F-3) | Santa Cruz | sc-55522 |
| PI 3-kinase p110 (D-4) | sc-8010 | sc-55522 |
| PCNA (PC10) | Santa Cruz | sc-56 |
| p-ERK (E-4) | Santa Cruz | sc-7383 |
| ERK | Bimake | A502 |
| p-AKT (Thr 308) | Santa Cruz | sc-16646. |
| p-EGFR (F3) | Santa Cruz | sc-377547 |
| KI67 | Santa cruz | sc-23900 |

| **­2^nd^ Antibodies** | **Company** | **Identifier** |
| --- | --- | --- |
| SuperBoost™ Goat anti-Mouse Poly HRP | ThermoFisher | B40961 |
| SuperBoost™ Goat anti-Rabbit Poly HRP | ThermoFisher | B40962 |
| Donkey anti-Mouse IgG (H+L) Highly Cross-Adsorbed Secondary Antibody, Alexa Fluor 488 | ThermoFisher | A21202 |
| Donkey anti-Rabbit IgG (H+L) Highly Cross-Adsorbed Secondary Antibody, Alexa Fluor 488 | ThermoFisher | A21206 |
| Donkey anti-Mouse IgG (H+L) Cross-Adsorbed Secondary Antibody, DyLight 680 | ThermoFisher | SA5-10170 |
| Donkey anti-Goat IgG (H+L) Cross-Adsorbed Secondary Antibody, DyLight 680 | ThermoFisher | SA5-10090 |
| Donkey anti-Rabbit IgG (H+L) Cross-Adsorbed Secondary Antibody, DyLight 800 | ThermoFisher | SA5-10044 |
| Donkey anti-Mouse IgG (H+L) Cross-Adsorbed Secondary Antibody, DyLight 800 | ThermoFisher | SA5-10172 |
| Donkey anti-Goat IgG (H+L) Cross-Adsorbed Secondary Antibody, DyLight 800 | ThermoFisher | SA5-10092 |

| **Bacterial Strains** | **Company** | **Identifier** |
| --- | --- | --- |
| **DH5α** F- endA1 glnV44 thi-1 recA1 relA1 gyrA96 deoR nupG Φ80dlacZΔM15 Δ(lacZYA-argF)U169, hsdR17(rK-mK+), λ– | ThermoFisher | 18263012 |
| **Chemicals and Commercial Assays** | **Company** | **Identifier** |
| Gibco™  Dulbecco’s Modified Eagle Medium (DMEM), high glucose | ThermoFisher | 11574486 |
| Gibco™ LHC Basal Medium (1X) | ThermoFisher | 11524556 |
| Gibco™ Trypsin-EDTA (0.5%), No Phenol Red | ThermoFisher | 15400054 |
| Fetal Bovine Serum (FCS) | Sigma-Aldrich | 12103C |
| Penicillin-Streptomycin | Sigma-Aldrich | P4333 |
| Polybrene | Sigma-Aldrich | TR-1003 |
| Polyethylenimine, Linear, MW 25000, Transfection Grade (PEI 25K) | Polysciences | 23966-1 |
| Dimethyl sulfoxide (DMSO) | Sigma-Aldrich | D8418 |
| Ethanol (Etoh) | Carl Roth | 5054.6 |
| Gibco™ Phosphate-buffered saline (PBS) | ThermoFisher | 10010031 |
| Nuclease-free water | Merck | 3098 |
| Pierce™ Protein A/G Magnetic Beads | ThermoFisher | 88802 |
| 2-Propanol/ Isopropanol | ROTH | AE73.2 |
| Adenosintriphosphat (ATP) | Jena Bioscience | NU-1010-10G |
| Agarose | ROTH | 3810.4 |
| Ampicillin (Amp) | ROTH | HP62.2 |
| Bovine serum albumine (BSA) | Merck Millipore | 810683 |
| Dithiothreitol (DTT) | Sigma-Aldrich | D9779 |
| Eosin | Sigma | E4009 |
| Hematoxylin | Sigma | H3136 |
| Polyvinylidene difluoride membranes (PVDF) Immobilon Transfer Membrane | Merck | IPFL00010 |
| N,N,N',N'-tetramethylenethylendiamine (TEMED) | ROTH | 2367.3 |
| Natrium chloride (NaCl) | AppliChem | A2942,1000 |
| Neutrally buffered formalin (NBF) | Thermo Fisher | 5700TS |
| Tris-HCl | ROTH | 9090.5 |
| TritonX100 | ROTH | 3051.3 |
| Xylene | Sigma | 534056 |
| β-Mercaptoethanol | ROTH | 4227.1 |
| Methanol (MeOH) | ROTH | 0082.3 |
| Random Hexamer Primer | ThermoFisher | SO142 |
| RiboLock RNase Inhibitor | ThermoFisher | EO0381 |
| Histo-Clear^®^ Histological Clearing Agent | National Diagnostics | HS-200; Lot03-19-18 |
| Cytoseal 60^™^ | Thermo Scientific | 8310-4 |
| SignalStainR DAB Substrate Kit | Cell Signaling | 8059 S |
| Doxycycline hyclate | Sigma-Aldrich | D9891 |
| AZ1 | Probechem | PC-60023 |
| AZ1 | Selleckhem | S8904 |
| Propidium Iodide (PI) | Sigma-Aldrich | 11348639001 |
| Vemurafenib | Selleckhem | S1267 |
| Buparlisib | Selleckhem | S2247 |
| Gefitinib | Selleckhem | S1025 |
| 4′,6-diamidino-2-phenylindole (DAPI) | ThermoFisher | D1306 |
| Hoechst | ThermoFisher | 62249 |
| **Experimental Models: Organisms/Strains Cell** | **Company** | **Identifier** |
| B6(C)-Gt(ROSA)26Sor^em1.1(CAG-cas9*,-EGFP)Rsky^/J | The Jackson laboratory | Stock No: 028555 |
| B6.129-Kras^tm4Tyj^ Trp53^tm1Brn^/J | The Jackson laboratory | Stock No: 032435 |
| C57BL/6J | The Jackson Laboratory | Stock No: 000664 |
| **Oligonucleotides and recombinant DNA** | **Sequence** | **Company** |
| shRNA hUSP28 #1 FW | CCGGCAAGGAGCTTATTCGAAATCTCGAGATTTCGAATAAGCTCCTTGTTTTTG | Sigma |
| shRNA hUSP28 #1 RV | AATTCAAAAACAAGGAGCTTATTCGAAATCTCGAGATTTCGAATAAGCTCCTTG | Sigma |
| shRNA hUSP28 #2 FW | CCGGGACTGAAGATCATCCATTACTCGAGTAATGGATGATCTTCAGTCTTTTTG | Sigma |
| shRNA hUSP28 #2 RV | AATTCAAAAAGACTGAAGATCATCCATTACTCGAGTAATGGATGATCTTCAGTC | Sigma |
| sgRNA mUsp28 #1 FW | CACCGGGGAGCCTTCCGATCATCCG | Sigma |
| sgRNA mUsp28 #1 RV | AAACCGGATGATCGGAAGGCTCCCc | Sigma |
| sgRNA mUsp28 #2 FW | CACCGCGGATCGTTCCGTGAAGTAT | Sigma |
| sgRNA mUsp28 #2 RV | AAACATACTTCACGGAACGATCCGc | Sigma |
| sgRNA mKras #1 FW | CACCGACTGAGTATAAACTTGTGG | Sigma |
| sgRNA mKras #1 RV | AAACCCACAAGTTTATACTCAGTC | Sigma |
| sgRNA mTrp53 #1 FW | CACCGATGGTGGTATACTCAGAGC | Sigma |
| sgRNA mTrp53 #1 RV | AAACGCTCTGAGTATACCACCATC | Sigma |
| mKrasG12D repair template FW | TTTTGTGTAAGCTTTGGTAACTCCATGTATTTTTATTAAGTGTT | Sigma |
| mKrasG12D repair template RV | GAGCTTATCGATACCGTCGACACACCCAGTTTAAAGCCTTGGAA | Sigma |
| pLKO.1 puro | pLKO.1 puro was a gift from Bob Weinberg (Addgene plasmid # 8453 ; http://n2t.net/addgene:8453) | Addgene plasmid # 8453 |
| pLKO shUSP_28_1 (Puromycin) | ^21^ | N/A |
| pLKO shUSP_28_2 (Puromycin) | ^21^ | N/A |
| pHelper | Cell Biolabs, INC. | VPK-400-DJ |
| pAAV-DJ Vector | Cell Biolabs, INC. | VPK-420-DJ |
| AAV:ITR- U6-sgRNA(p53)- pEFS-2A-mCherry-shortPA- ITR | ^21^ | N/A |
| AAV:ITR-U6-sgRNA(Kras)-pEFS-2A-mCherry-shortPA-KrasG12D_HDRdonor-ITR | ^21^ | N/A |
| AAV:ITR-U6-sgRNA(Kras)-U6-sgRNA(p53)-pEFS-2A-mCherry-shortPA-KrasG12D_HDRdonor-ITR | ^21^ | N/A |
| pHelper | Cell Biolabs, INC. | VPK-400-DJ |
| pAAV-DJ Vector | Cell Biolabs, INC. | VPK-420-DJ |
| pUMVC | pUMVC was a gift from Bob Weinberg | Addgene plasmid # 8449 |
| pCMV-VSV-G | pCMV-VSV-G was a gift from Bob Weinberg | Addgene plasmid # 8454 |
| EGFR | EGFR was a gift from Matthew Meyerson (Greulich H, Chen TH et al. 2005). | Addgene number: #11011 |
| EGFR L858R | EGFR L858R was a gift from Matthew Meyerson (Greulich H, Chen TH et al. 2005). | Addgene number: #11012 |
| pBabe puro HA PIK3CA | pBabe puro HA PIK3CA were a gift from Jean Zhao (Zhao JJ, Liu Z, Wang L et al. 2005) | addgene number: #12522 |
| pBabe puro HA PIK3CA H1047R | pBabe puro HA PIK3CA H1047R were a gift from Jean Zhao (Zhao JJ, Liu Z, Wang L et al. 2005) | addgene number: #12524 |
| pBabe puro HA PIK3CA E545K | pBabe puro HA PIK3CA E545K were a gift from Jean Zhao (Zhao JJ, Liu Z, Wang L et al. 2005) | addgene number: #12525 |
| pBabe puro HA PIK3CA | pBabe puro HA PIK3CA were a gift from Jean Zhao (Zhao JJ, Liu Z, Wang L et al. 2005) | addgene number: #12522 |
| pBabe puro HRAS G12D | This Manuscript | NA |
| pBabe Puro BRAF V600E | pBabe Puro BRAF V600E was a gift from William Hah | addgene number: # 15269 |

| **Software** | **Company/Source** |
| --- | --- |
| cBioportal | https://www.cbioportal.org |
| GEPIA and GEPIA2 | http://gepia.cancer-pku.cn |
| KM-plotter | http://kmplot.com/analysis/ |
| Operetta Imaging | Perkin Elmer |
| BoxPlotR | http://shiny.chemgrid.org/boxplotr/ |
| Morpheus | https://software.broadinstitute.org/morpheus/ |
| Excel | Microsoft |
| Affinity Desgner | h https://affinity.serif.com/es/designer/ |
| Image Studio | Licor |
| Panther Classification system | http://pantherdb.org |
| AATBIO IC50 calculator | https://www.aatbio.com/tools/ic50-calculator |
| PRISM4 | GraphPad Software, Inc. |
| Affinity Designer | Serif Europe |
| ImageJ | National Insistute of Health |
| Primerx | http://www.bioinformatics.org/primerx/cgi-bin/DNA_1.cgi |
| ROC Plotter | http://www.rocplot.org/ |
| Pannoramic Case Viewer | 3dHistech |
| R2: Genomics Analysis and Visualization Platform | http://r2.amc.nl |
| UCSC Xena | https://ucsc-xena.gitbook.io/project/ |
| Proteome discoverer 2.2 | Thermo Scientific |
| MaxQuant | https://www.maxquant.org/ |
| Perseus 1.6.5. | https://maxquant.net/perseus/ |
| Uniprot | https://www.uniprot.org/ |
| INSTANT CLUE | https://kups.ub.uni-koeln.de/17610/ |
| FastQC | http://www.bioinformatics.babraham.ac.uk/projects/fastqc/ |
| Bowtie2 v2.3.4.1 | http://bowtie-bio.sourceforge.net/index.shtml |
| TopHat v.2.1.1 | <https://ccb.jhu.edu/software/tophat/index.shtml> |
| Samtools v1.3 | http://samtools.sourceforge.net |
| R | https://www.r-project.org |
| EdgeR | <https://bioconductor.org/packages/release/bioc/html/edgeR.html> |
| COMBENEFIT | <https://www.cruk.cam.ac.uk/research-groups/jodrell-group/combenefit> |
| Harmony Software | Perkin Elmer |
| EMBL | <https://www.embl.de/> |
| SPLASHRNA | <http://splashrna.mskcc.org/> |
| BD FACSDiva 6.1.2 BD | Biosciences |
| FlowJo 8.8.6 | FlowJo, LLC |
| Illustrator TM, | Adobe Inc. |
| Photoshop TM, | Adobe Inc. |
| Acrobat TM | Adobe Inc. |
| Integrated Genome Browser | Nicol et al. 2009 |
| Mac OS X | Apple Inc. |
| Office 2011 Mac | Microsoft Inc. |
| Qupath | https://qupath.github.io |
| UCSC Genome Bioinformatics | http://genome.ucsc.edu |
| ApE plasmid editor | By Wayne Davis |
| Venn diagrams | http://bioinformatics.psb.ugent.be/webtools/Venn/ |
| Nemates | http://nemates.org |
| Online Web statistical calculator Astatsa | https://astatsa.com/ |
| DOI citation formatter | https://citation.crosscite.org |
| Zhang lab gRNAs design resources | https://zlab.bio/guide-design-resources |
| CHOPCHOP | http://chopchop.cbu.uib.no/ |
| RNAi Consortium | www.broadinstitute.org/rnai-consortium/rnai-consortium-shrna-library |
| Cancer Therapeutics Response Portal | https://portals.broadinstitute.org/ctrp.v2.1/ |
| Genomics of Drug Sensitivity in Cancer | https://www.cancerrxgene.org/ |
| Catalogue Of Somatic Mutations In Cancer (COSMIC) | https://cancer.sanger.ac.uk/cell_lines |
| Gene Expression and Mutations in Cancer Cell Lines (GEMiCCL) | https://www.kobic.kr/GEMICCL/ |
| Clust Vis | https://biit.cs.ut.ee/clustvis/ |
| Depmap | https://depmap.org/portal/ |
| **Instrument** | **Company** |
| Odyssey® CLx Imaging System | Licor |
| iBright™ FL1000 Imaging System | Invitrogen |
| BD FACSCanto II Cell Analyzer | BD Biosciences |
| StepOnePlus Real-Time PCR System | Thermo Scientific |
| Invitrogen Countess II FL Automated Cell Counter | Thermo Scientific |
| Pannoramic DESK scanner | 3DHISTECH |
| FSX100 microscopy | Olympus Life Science |
| Operetta High-Content Imaging System | Perkin Elmer |
| Fragment Analyzer | Agilent formerly Advanced Analytical |
| Axiocam 503 mono | Zeiss |
| Branson Sonifier 250 | Branson |
| 250 mm long C18 column: X-Bridge, 4.6 mm ID, 3.5 µm particle size | Waters |
| EASY-nLC™ 1200 System | Thermo Scientific |
| Orbitrap Fusion Lumos mass spectrometer | Thermo Scientific |
| 1.9 µm C18 particles | ReproSil-Pur, Dr. Maisch |
| Hyrax M55 Rotary Microtome | Leica |
| Mr. Frosty freezer container | Thermo Scientific |
| PCR cycler: SimpliAmp thermo cycler | Life technologies |
| Experion™ Automated Electrophoresis  System | Bio-Rad |
| Cell culture incubator BBD 6220 | Heraeus |
| Casy® cell counter | Innovatis |
| Centrifuge Avanti J-26 XP | Backman Coulter |
| Centrifuge Eppendorf 5417 R | Eppendorf |
| Centrifuge Eppendorf 5425 | Eppendorf |
| Centrifuge Eppendorf 5430 | Eppendorf |
| Centrifuge Galaxy MiniStar | VWR |
| Centrifuge Multifuge 1S-R | Heraeus |
| Deep-sequencer Genome Analyzer IIx | Illumina |
| Dry Bath System | Starlab |
| Thermomixer® comfort | Eppendorf |
| Incubator shaker Model G25 | New Brunswick Scientific |
| Luminometer GloMax | Promega |
| Microscopes Axiovert 40CFL | Zeiss |
| PCR thermal cycler Mastercycler pro S | Eppendorf |
| Spectrofluorometer NanoDrop 1000 | Thermo Scientific |
| UltrospecTM 3100 pro UV/Visible | Amersham Biosciences |
| SDS page system Minigel | Bio-Rad |
| SDS page system Tetra Cell | Bio-Rad |
| Maxi UV fluorescent table | Peqlab |
| Mixer Vortex-Genie 2 | Scientific Industries |
| Julabo ED-5M water bath | Julabo |
| Memmert waterbath | Memmert |
| Immunoblot transfer system: Perfect Blue Tank Electro Blotter Web S | Peqlab |
| Power supply: Power Pac | Bio-Rad |
| Chemiluminescence imaging LAS-4000 mini Fujifim | Fujifim |
| Sterile bench HeraSafe | Heraeus |
| Siemens linear accelerator for X-ray irradiation | Siemens |
| Pipetman Classic P2.5, P10, P20, P200 and P1000 | Gilson |
| Dionex Ultimate 3000 analytical HPLC | Thermo Scientific |
| Leica VT 1200S | Leica |
| Microscope TCS SP5 | Leica |
| BD FACS Aria III | BD Biosciences |
| Pipetboy acu 2 | Integra |
| Consort EV243 electrophoresis power supply | Sigma |
| Ventana DP 200 slide scanner | Roche |

| **Cell lines**  **Cell** | **Source** | **Identifier** |
| --- | --- | --- |
| Human: HEK 293T | ATCC | CRL-11268 |
| BEAS-2B UD | Cell line was a gift from Marco A. Calzado Canale | CRL-9609 |
| BEAS-2B DIF | Cell line was a gift from Marco A. Calzado Canale | CRL-9609 |
| BEAS-2B ONC EGFR L858R | This Manuscript | NA |
| BEAS-2B ONC PIK3CA | This Manuscript | NA |
| BEAS-2B ONC PIK3CA H1047R | This Manuscript | NA |
| BEAS-2B ONC PIK3CA E545A | This Manuscript | NA |
| BEAS-2B ONC BRAF V600E | This Manuscript | NA |
| BEAS-2B ONC HRAS G12D | This Manuscript | NA |
| **RT-PCR primers** | **Sequence** | **Company** |
| RT-PCR hUsp28 FW | ACTCAGACTATTGAACAGATGTACTGC | Sigma |
| RT-PCR hUsp28 RV | CTGCATGCAAGCGATAAGG | Sigma |
| RT-PCR hB-Actin FW | GCTACGAGCTGCCTGACG | Sigma |
| RT-PCR hB-Actin RV | GGCTGGAAGAGTGCCTCA | Sigma |
| RT-PCR hTp63 FW | GGAAAACAATGCCCAGACTC | Sigma |
| RT-PCR hTp63 RV | GTGGAATACGTCCAGGTGGC | Sigma |
| RT-PCR hKeratin14 FW | GGAAGTGAAGATCCGTGACTG | Sigma |
| RT-PCR hKeratin14 RV | GGACTGTAGTCTTTGATCTCAGCA | Sigma |
| RT-PCR mB-Actin FW | AGTGTGACGTTGACATCCGT | Sigma |
| RT-PCR mB-Actin RV | TGCTAGGAGCCAGAGCAGTA | Sigma |
| RT-PCR mUSP28 FW | ATGACAACTTGCCCCACTTC | Sigma |
| RT-PCR mUSP28 RV | AGTTCCACAGACAGGGCTTC | Sigma |
| RT-PCR hEgfr FW | AACACCCTGGTCTGGAAGTACG | Sigma |
| RT-PCR hEgfr RV | TCGTTGGACAGCCTTCAAGACC | Sigma |
| RT-PCR hHras FW | ACGCACTGTGGAATCTCGGCAG | Sigma |
| RT-PCR hHras RV | TCACGCACCAACGTGTAGAAGG | Sigma |
| RT-PCR hBraf FW | AACGAGACCGATCCTCATCAGC | Sigma |
| RT-PCR hBraf RV | GGTAGCAGACAAACCTGTGGTTG | Sigma |
| RT-PCR hPik3CA FW | GAAGCACCTGAATAGGCAAGTCG | Sigma |
| RT-PCR hPik3CA RV | GAGCATCCATGAAATCTGGTCGC | Sigma |
