## Supplementary material for "USP28 enables oncogenic transformation of respiratory cells and its inhibition potentiates molecular therapy targeting mutant EGFR, BRAF and PI3K": Suppl. Figure legends 1-7

### Supplementary Figure Legends

#### Figure S1: *USP28 is upregulated irrespective of lung tumour subtype*

A) Publicly available data of relative mRNA expression of USP28 in human NSCLC Adenocarcinoma samples, separated by subclass. Shown are non-transformed and tumour samples. N = normal lung tissue, T = tumour. n indicates number of individual samples. Generated with the open-source tool www.gepia2.cancer-pku.cn. p-values were calculated using two-tailed T-test statistical analysis. p<0.005 = ***. In box plots, the centre line reflects the median and the upper and lower box limits indicate the first and third quartiles. Whiskers extend 1.5× the IQR.

B) Publicly available data of relative mRNA expression of USP28 in human NSCLC Squamous cell carcinoma samples, separated by subclass. Shown are non-transformed and tumour samples. N = normal lung tissue, T = tumour. n indicates number of individual samples. Generated with the open-source tool www.gepia2.cancer-pku.cn. p-values were calculated using two-tailed T-test statistical analysis. p<0.0005 = ****. In box plots, the centre line reflects the median and the upper and lower box limits indicate the first and third quartiles. Whiskers extend 1.5× the IQR.

#### Figure S2: *USP28 is required to establish oncogenic transformation in vivo.*

A) Overview of genetic alterations within the ERBB, RAS, PI3K and RAF oncogenic families in NSCLC (SCC and ADC). Shown are total percent of genetic alterations within the respective pathway and individual pathway members. N = 501 SCC and n = 515 ADC samples. Data generated with the online tools www.cbioportal.org.

B) *Schematic representation of in vivo* CRISPR mediated gene editing to induce deletion of *Trp53* and mutation of *Braf* to *Braf*$^{V600E}$ (PB) upon intratracheal instillation of AAV virions, packaged with the pleotropic capsid 2/DJ, into *C57Bl6/J-Rosa26*$^{Sor\text{-}CAGG\text{-}Cas9\text{-}IRES\text{-}eGFP}$ mice. Animals were sacrificed 4 weeks post infection. n= 3.

C) Representative images of immuno-histochemical staining against endogenous Usp28, Nkx2-1/Ttf-1 and the proliferative marker Pcna in murine lungs infected with either control virus (WT) or PB. Shown a representative lung, with a low and high magnification of individual areas. Animals were analysed 4 weeks' post-infection n=3

#### Figure S3: *Malignant transformation of BEAS-2B*$^{DIF}$ *via EGFR-PI3K-MAP.*

A) mRNA expression correlation data between USP28 and RAS family members H-RAS, K-RAS and NRAS in NSCLC ADC and SCCC, obtained by public available data sets (www.gepia2.cancer-pku.cn/ ). Shown is the Spearman correlation efficiency (R).

B) Graphical visualization of the Spearman´s correlation between USP28 and KRAS or NRAS in ADC tumours. Correlation and p-value calculated were calculating using the online tool GEPIA www.gepia2.cancer-pku.cn/ .

C) Schematic overviews of commonly occurring mutations in NSCLC driver oncogenes EGFR, HRAS, BRAF and PIK3CA. Shown are missense (green), truncating mutations (black), in frame mutations (brown) and other genetic alterations (violet). Data was generated using the online tool www.cbioportal.org.
D) Immunoblot against EGFR, HRAS, BRAF and PIK3CA oncogenic drivers in BEAS-$2B^{DIF}$ or BEAS-$2B^{DIF}$ upon retroviral transduction to express the indicated oncogenes EGFR (wild type (WT) and L858R), HRAS (G12D), BRAF (V600E) and PIK3CA (wild type (WT), E545K and H1047R), respectively. ACTIN served as loading control. Representative immunoblot of n=3.
E) RT-PCR of EGFR, HRAS, BRAF and PIK3CA oncogenic drivers in BEAS-$2B^{DIF}$ or BEAS-$2B^{DIF}$ upon retroviral transduction to express the indicated oncogenes EGFR (wild type (WT) and L858R), HRAS (G12D), BRAF (V600E) and PIK3CA (wild type (WT), E545K and H1047R), respectively. Shown are log2 fold change expression data, relative to ACTIN and normalized to the respective expression in BEAS-$2B^{DIF}$. Shown are mean values and standard deviation of n=3

**Figure S4:** ***Oncogenic transformation via EGFR-PI3K-MAPK pathway upregulate USP28.***

A) Schematic model. c-JUN and c-MYC up-regulate transcription of USP28 (Serra et al. 2014; Diefenbacher et al. 2014).
B) Immunoblot against USP28, c-MYC, c-JUN in BEAS-$2B^{DIF}$ or BEAS-$2B^{DIF}$ upon transfection of cDNA encoding c-JUN and c-MYC. VINCULIN served as loading control. Representative immunoblot of n=3.
C) Immunoblot against endogenous USP28 and its substrates NOTCH1, c-MYC and c-JUN of BEAS-$2B^{UD}$, BEAS-$2B^{DIF}$ and BEAS-$2B^{DIF}$ upon retroviral transduction to express the indicated oncogenes EGFR (L858R), HRAS (G12D), BRAF (V600E) and PIK3CA (wild type (WT) and H1047R), respectively. VINCULIN or ACTIN served as loading control. Representative immunoblot of n=3.
D) Overview of the oncogenic pathways and recurring oncogenic drivers involved in the regulation of USP28 expression and abundance. Oncogenic transformation upon activation of EGFR-PI3K-MAPK pathway up-regulates transcription of USP28 via c-MYC/ c-JUN.
A) Pan-Cancer mRNA expression correlation data between USP28 and AKT2 or BRAF, obtained by public available data sets (Depmap.org). Pearson´s correlation between USP28 and AKT2 or BRAF mRNA levels in cancer cell lines from different tumor entities. For graphical visualization, linear regression was included and every dot represent a different cell line. The table indicates the next parameters for different tumor entities: Pearson correlation, slope and p-value for the individual tumour types, relative to AKT2-USP28 and BRAF-USP28. The parameters were calculated by the online tool https://depmap.org/.

**Figure S5:** ***Malignant transformation renders tumour cells dependent on USP28***

A) Immunofluorescence of USP28 endogenous expression in A549 cells. A549 is a KRAS mutated cell line (KRASG12S). ACTIN and DAPI as control markers.

B) Immunoblot showing protein abundance of USP28, c-MYC and PCNA in A549 upon lentiviral transduction with either a control or two individual constitutive shRNA targeting USP28. ACTIN served as loading control.

C) For growth analysis, A549 cells from B) were seeded at equal cell density and counted at day 1, day 3 and day 5. p-values were calculated using two-tailed T-test statistical analysis. Shown are mean values and standard deviation of n=3.

D) BEAS-2B$^{ONC}$ (EGFRL858R, PIK3CA L1047R and BRAFV600E) cultured in the presence of either control solvent (DMSO) or AZ1 for 72 hours at indicated concentrations. Shown are representative DAPI Images of cells 72 hours post culture in the presence of DMSO or AZ1. Calculation of GI50 was performed upon cell quantification of 30-45 20x fields from independent wells in control (DMSO) and AZ1 conditions.

Figure S6: ***Inhibition of USP28 via AZ1 'resets' the proteome of oncogenic transduced cells towards a 'non-oncogenic' state and induces pro-apoptotic signatures***

A) Schematic model explaining the proteomics experiments and analysis performed in this manuscript.

B) Volcano plots comparing protein abundances in BEAS-2B$^{DIF}$ and BEAS-2B$^{ONC}$ (EGFRL858R, PIK3CA L1047R and BRAFV600E).

C) Volcano plots comparing protein abundances in BEAS-2B$^{ONC}$ (EGFRL858R, PIK3CA L1047R and BRAFV600E) upon exposure to DMSO or 15 µM AZ1 for 72 hours AZ1.

D) Heatmap of proteins identified in Figure 6B for BEAS-2B$^{DIF}$, BEAS-2B$^{ONC}$ (PIK3CA L1047R or BRAFV600E) and BEAS-2B$^{ONC}$ (PIK3CA L1047R or BRAFV600E) treated with 15 µM AZ1 for 72 hours. Shown are n=3 experiments and data presented as Z score values per row. Red= high Z-score, blue = low Z-score protein abundance.

Figure S7: ***USP28 inhibition potentiates targeted molecular therapy***

A) Schematic model explaining the concept of overall survival (OS), progression free survival/disease (PFS) and post-progression survival (PPS).

B) Kaplan-Meier plots of NSCLC patient post-progression survival (PPS), relative to USP28 expression for total number of patients, patients at Stage 1 and patients at Stage 2. Data was generated using the online tool www.kmplot.com.

C) Analysis of publicly available datasets analysing USP28 mRNA expression and Overall Survival (OS) in NSCLC. Samples were divided in two groups based on USP28 mRNA expression: High USP28 (higher than the median USP28 expression) and Low USP28 (lower than the median USP28 expression). Survival days was determined for both groups. Data was obtained from the online tool https://xena.ucsc.edu/. In box plots, the centre line reflects the median and the upper and lower box limits indicate the first and third quartiles. Whiskers extend 1.5× the IQR.

D) Immunoblot showing protein abundance of p-EGFR, p-AKT, P-ERK and ERK in BEAS-2B$^{ONC}$ (EGFRL858R, PIK3CA L1047R and BRAFV600E) upon exposure to DMSO or 20 μM Gefitinib/ 20 μM Vemurafenib/ 1 μM Buparlisib for 72 hours. VINCULIN serves as loading control. n=3

E) BEAS-2B$^{ONC}$ (EGFRL858R, PIK3CA L1047R and BRAFV600E) cultured in the presence of either control solvent (DMSO) 15 μM AZ1 or 20 μM Gefitinib/ 20 μM Vemurafenib/ 0.5 μM Buparlisib for 144 hours at indicated concentrations. Viability was quantified upon Crystal violet staining of 3 independent wells Shown are crystal violet representative stained wells. Shown are mean values and standard deviation. p-values were calculated using two-tailed T-test statistical analysis.
