## Supplementary material for "USP28 enables oncogenic transformation of respiratory cells and its inhibition potentiates molecular therapy targeting mutant EGFR, BRAF and PI3K": Suppl. Figure 1-7

A

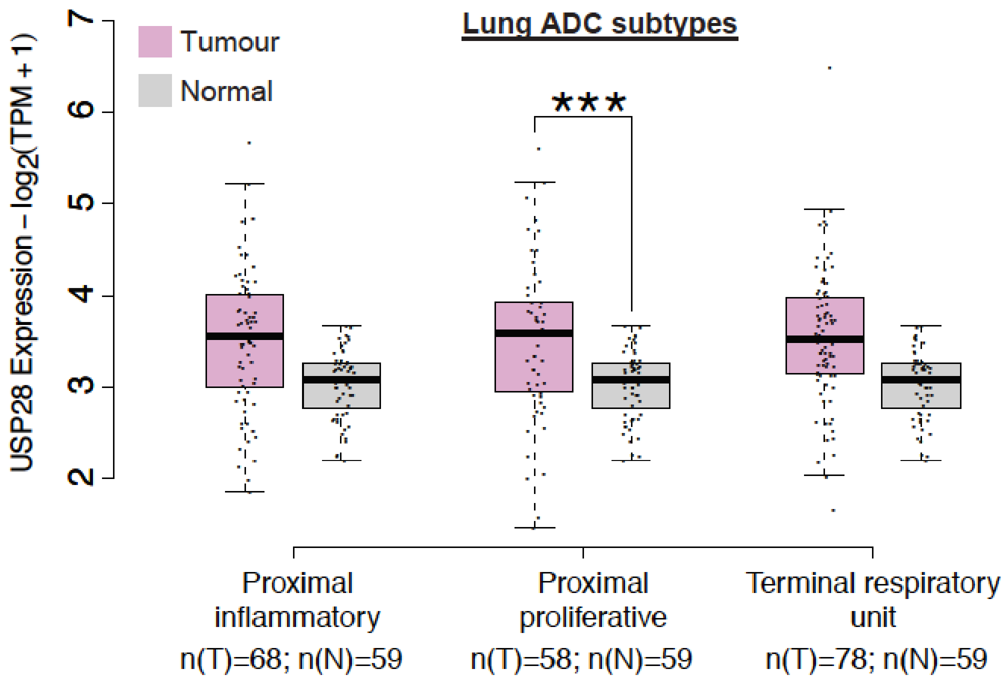


B

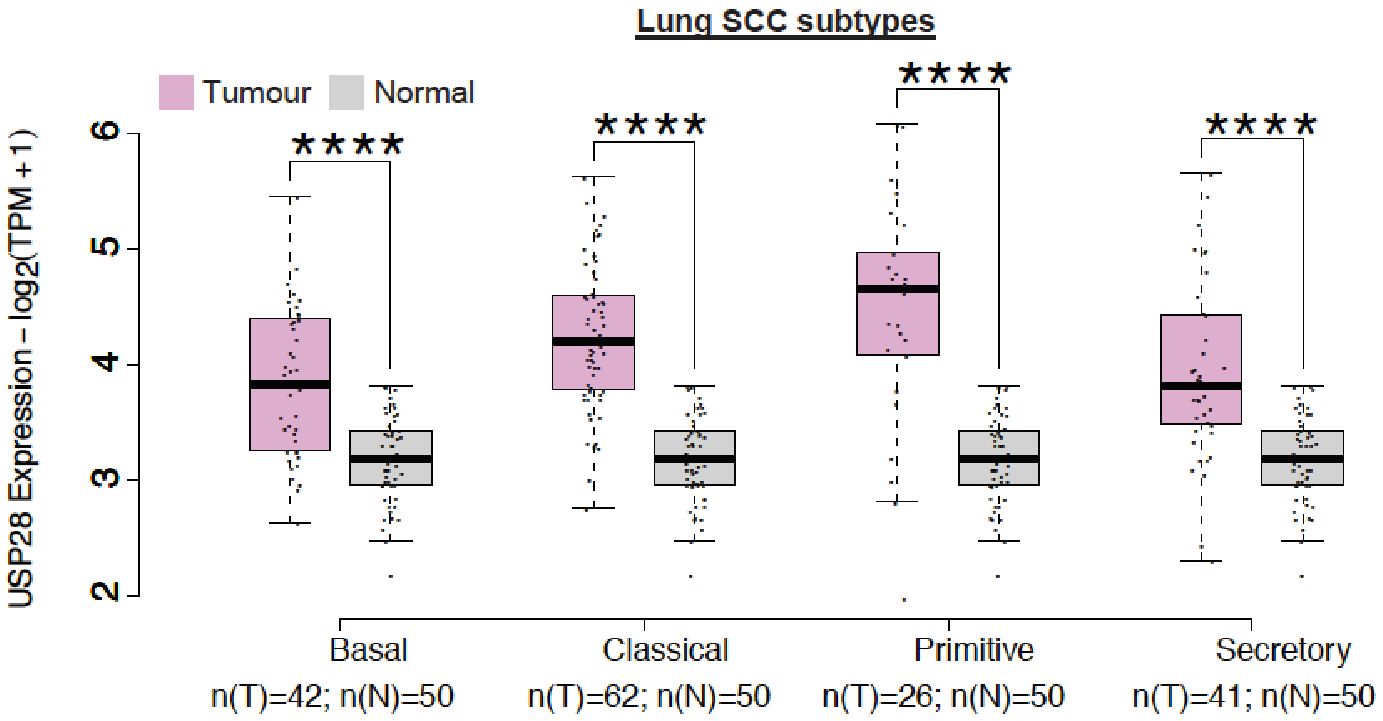


Prieto-Garcia et al. Figure S1

A

| | ErbB Family | | RAS Family | | PIK3 class1 Family | | RAF Family | |
|---|---|---|---|---|---|---|---|---|
| N=501 | SCC (43% altered) | | SCC (45% altered) | | SCC (85% altered) | | SCC (63% altered) | |
| | EGFR | 13% | HRAS | 8% | PIK3CA | 77% | A-RAF | 19% |
| | HER1 | 17% | **KRAS** | 25% | PIK3CB | 35% | **B-RAF** | 28% |
| | HER2 | 13% | NRAS | 22% | PIK3CD | 3% | C-RAF | 39% |
| | HER3 | 13% | | | PIK3CG | 11% | | |
| N=515 | ADC (59% altered) | | ADC (47% altered) | | ADC (50% altered) | | ADC (51% altered) | |
| | EGFR | 28% | HRAS | 8% | PIK3CA | 29% | A-RAF | 14% |
| | HER1 | 26% | **KRAS** | 27% | PIK3CB | 18% | **B-RAF** | 25% |
| | HER2 | 20% | NRAS | 22% | PIK3CD | 12% | C-RAF | 28% |
| | HER3 | 9% | | | PIK3CG | 13% | C-RAF= Down-reg. | |

B

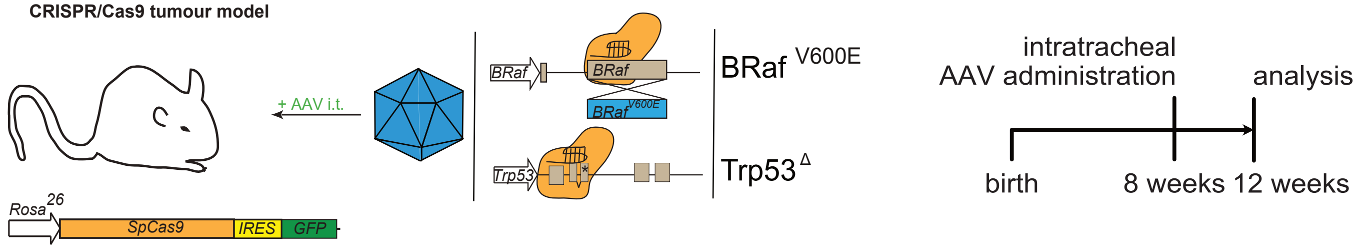


C

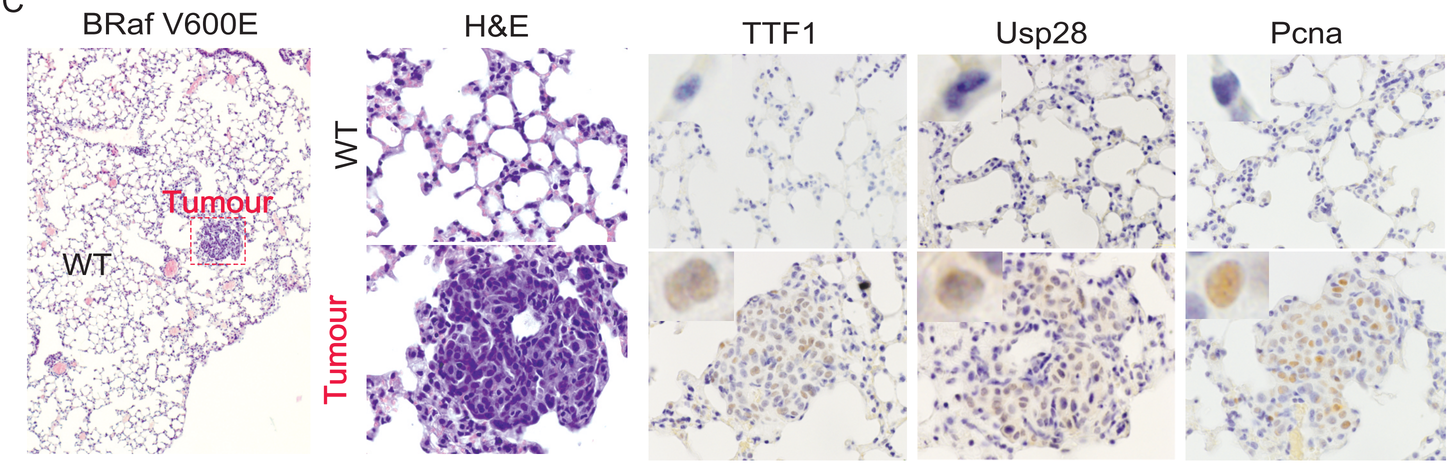

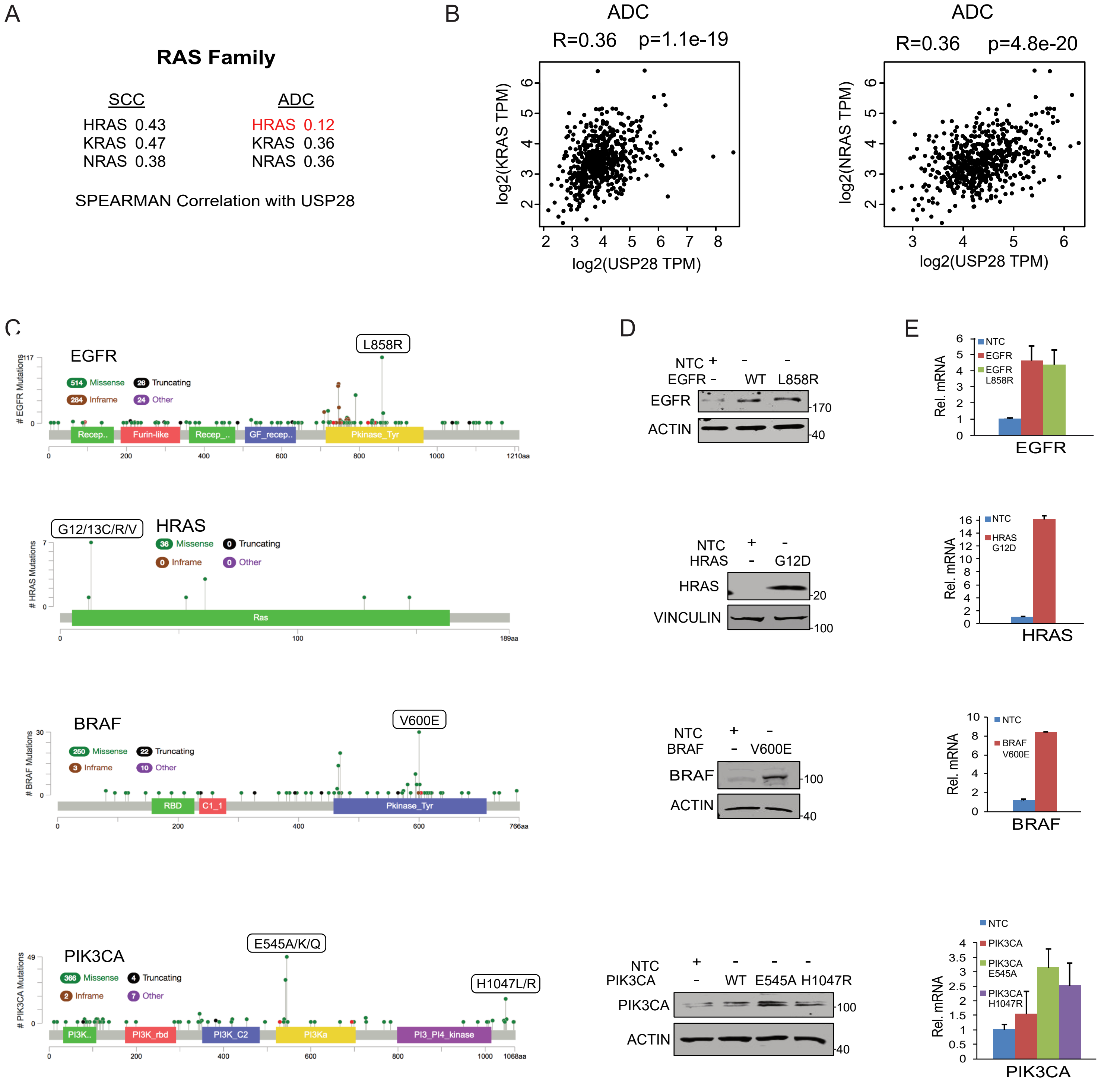


Prieto-Garcia et al. Figure S3

A

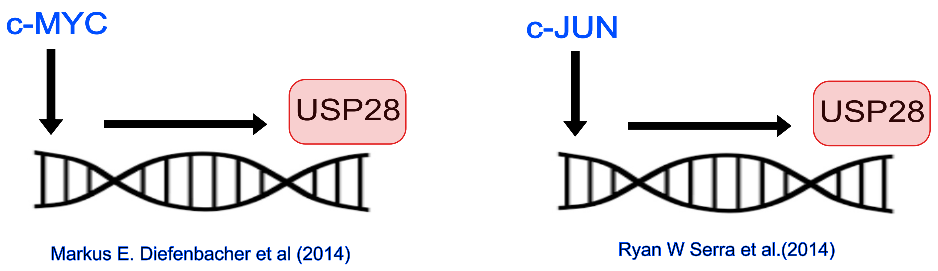


B

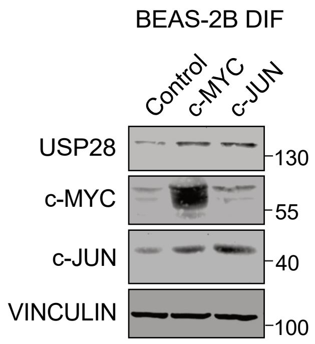


C

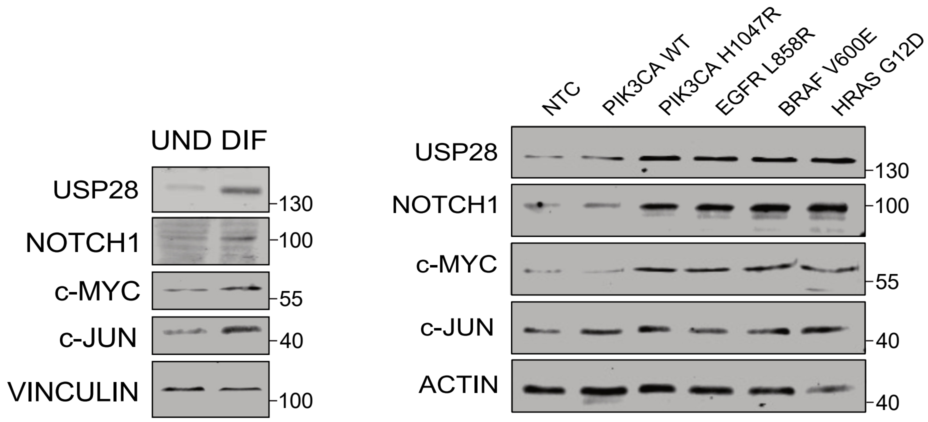


D

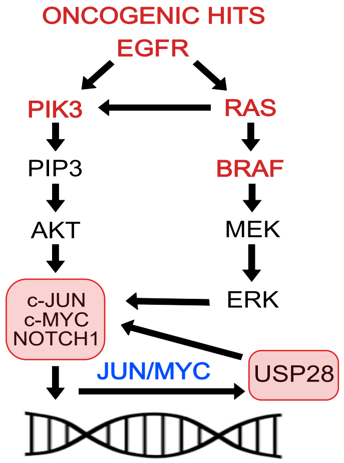


E

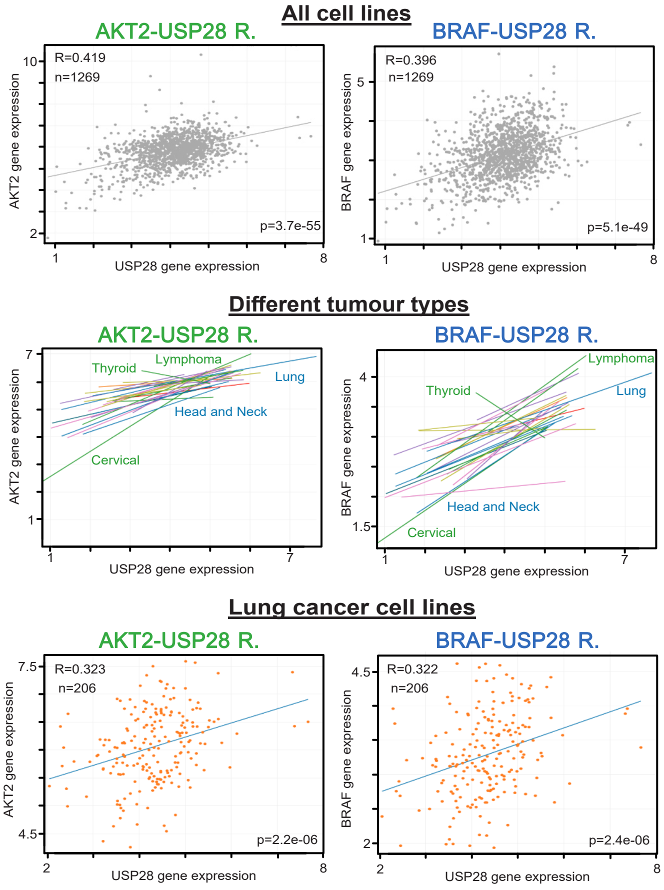


| Cell line Tumour type | AKT2-USP28 R. Pearson R | AKT2-USP28 R. Slope | AKT2-USP28 R. p-value | BRAF-USP28 R. Pearson R | BRAF-USP28 R. Slope | BRAF-USP28 R. p-value |
|---|---|---|---|---|---|---|
| Thyroid Cancer | -0.253 | -2.72E-1 | 4.28E-1 | -0.665 | -4.40E-1 | 1.82E-2 |
| Skin Cancer | 0.244 | 2.04E-1 | 4.34E-2 | 0.090 | 7.03E-2 | 4.62E-1 |
| Sarcoma | 0.579 | 4.16E-1 | 2.72E-4 | 0.386 | 2.60E-1 | 2.22E-2 |
| Rhabdoid | 0.569 | 8.10E-1 | 8.78E-3 | 0.467 | 6.02E-1 | 3.78E-2 |
| Prostate Cancer | 0.300 | 2.59E-1 | 5.13E-1 | 0.580 | 4.75E-1 | 1.72E-1 |
| Pancreatic Cancer | 0.427 | 5.44E-1 | 1.99E-3 | 0.422 | 2.68E-1 | 2.29E-3 |
| Ovarian Cancer | 0.321 | 3.50E-1 | 1.50E-2 | 0.569 | 4.00E-1 | 3.91E-6 |
| Neuroblastoma | 0.550 | 4.60E-1 | 3.59E-3 | 0.342 | 3.42E-1 | 8.74E-2 |
| Myeloma | 0.112 | 8.42E-2 | 5.56E-1 | 0.270 | 3.09E-1 | 1.48E-1 |
| Lymphoma | 0.685 | 6.06E-1 | 9.21E-12 | 0.602 | 4.84E-1 | 8.67E-9 |
| Lung Cancer | 0.323 | 2.54E-1 | 2.24E-6 | 0.322 | 2.34E-1 | 2.38E-6 |
| Liver Cancer | 0.068 | 4.60E-2 | 7.51E-1 | 0.110 | 8.65E-2 | 6.10E-1 |
| Liposarcoma | 0.818 | 5.94E-1 | 7.00E-3 | 0.218 | 6.46E-2 | 5.73E-1 |
| Leukemia | 0.436 | 3.09E-1 | 1.14E-5 | 0.378 | 2.45E-1 | 1.72E-4 |
| Kidney Cancer | 0.418 | 2.79E-1 | 1.55E-2 | 0.481 | 3.59E-1 | 4.61E-3 |
| Head and Neck Cancer | 0.580 | 4.91E-1 | 8.86E-5 | 0.573 | 4.40E-1 | 1.11E-4 |
| Gastric Cancer | 0.364 | 4.66E-1 | 1.95E-2 | 0.334 | 3.28E-1 | 3.30E-2 |
| Gallbladder Cancer | 0.834 | 4.60E-1 | 7.89E-2 | 0.480 | 1.71E-1 | 4.13E-1 |
| Fibroblast | 0.428 | 3.06E-1 | 6.60E-3 | 0.269 | 1.63E-1 | 9.75E-2 |
| Eye Cancer | 0.120 | 6.63E-2 | 7.98E-1 | 0.502 | 2.90E-1 | 2.51E-1 |
| Esophageal Cancer | 0.385 | 3.43E-1 | 2.98E-2 | 0.409 | 2.79E-1 | 2.02E-2 |
| Endometrial/Uterine Cancer | 0.303 | 3.99E-1 | 9.19E-2 | 0.672 | 5.28E-1 | 2.51E-5 |
| Colon/Colorectal Cancer | 0.575 | 4.52E-1 | 4.41E-7 | 0.521 | 3.37E-1 | 7.38E-6 |
| Cervical Cancer | 0.935 | 9.08E-1 | 8.25E-6 | 0.760 | 4.59E-1 | 4.12E-3 |
| Breast Cancer | 0.336 | 3.53E-1 | 1.15E-2 | 0.338 | 2.70E-1 | 1.08E-2 |
| Brain Cancer | 0.244 | 1.69E-1 | 3.12E-2 | 0.009 | 5.47E-3 | 9.35E-1 |
| Bone Cancer | 0.107 | 9.25E-2 | 5.36E-1 | 0.342 | 3.08E-1 | 4.09E-2 |
| Bladder Cancer | 0.191 | 1.72E-1 | 2.63E-1 | 0.212 | 1.66E-1 | 2.15E-1 |
| Bile Duct Cancer | 0.416 | 3.31E-1 | 1.78E-2 | 0.449 | 2.65E-1 | 9.92E-3 |

Prieto-Garcia et al. Figure S4

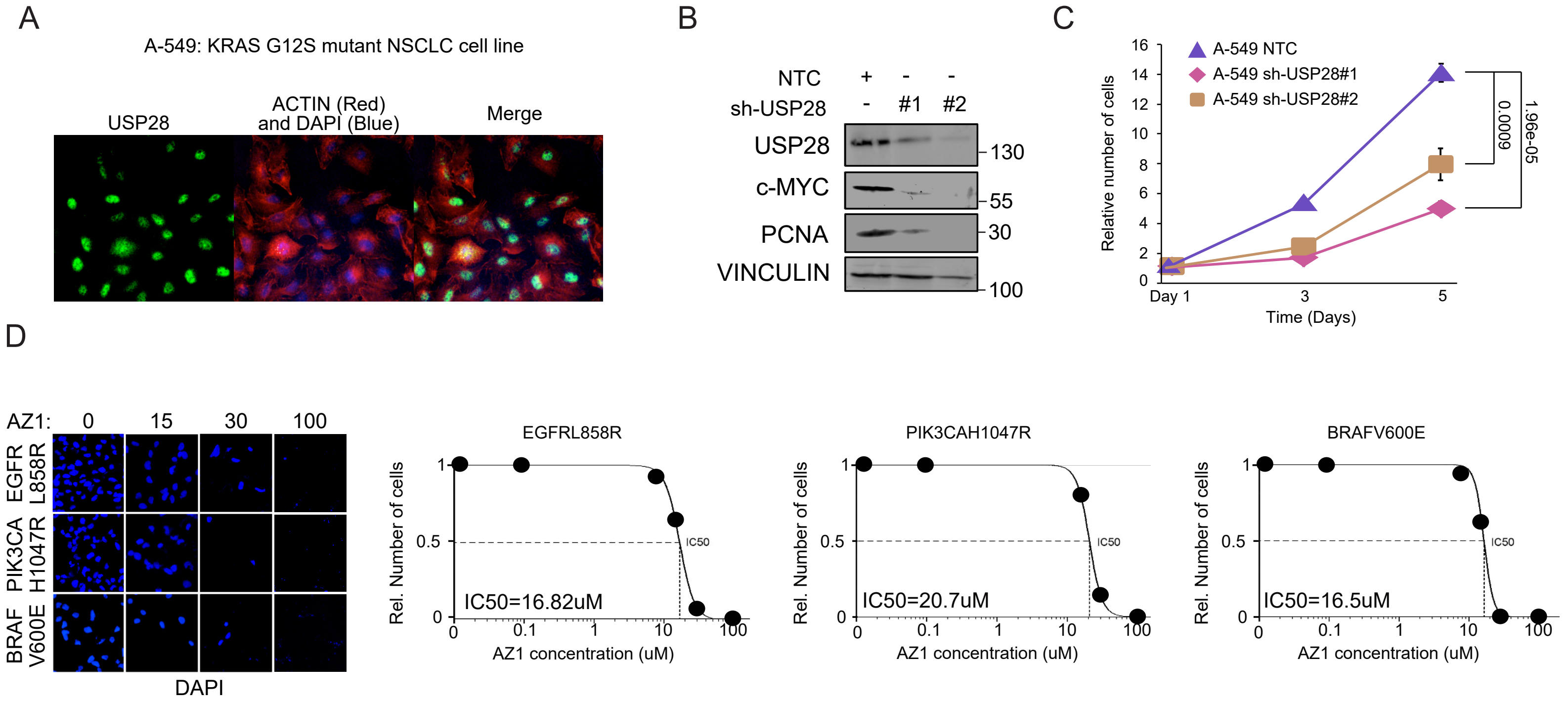


Prieto-Garcia et al. Figure S5

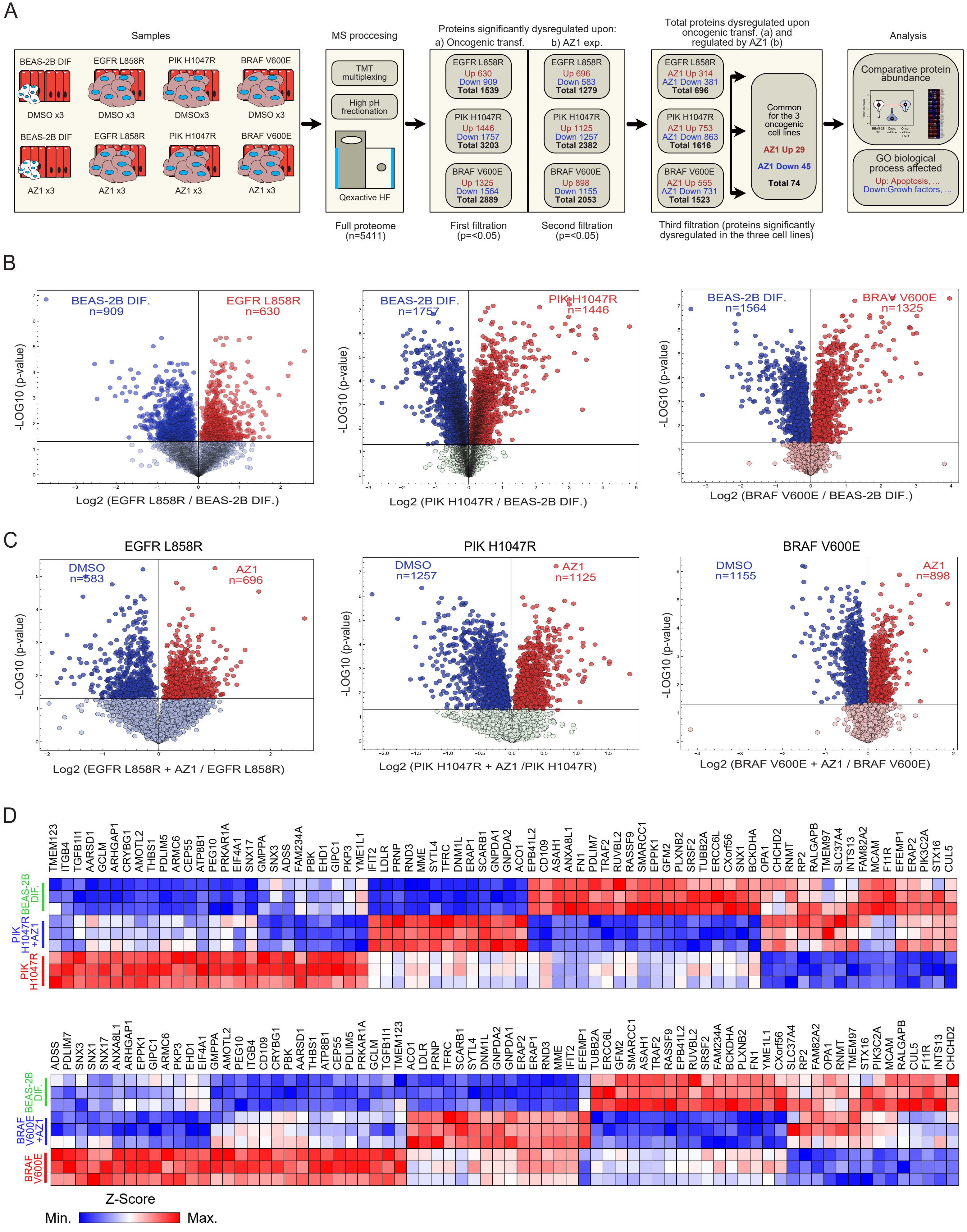


Prieto-Garcia et al. Figure S6

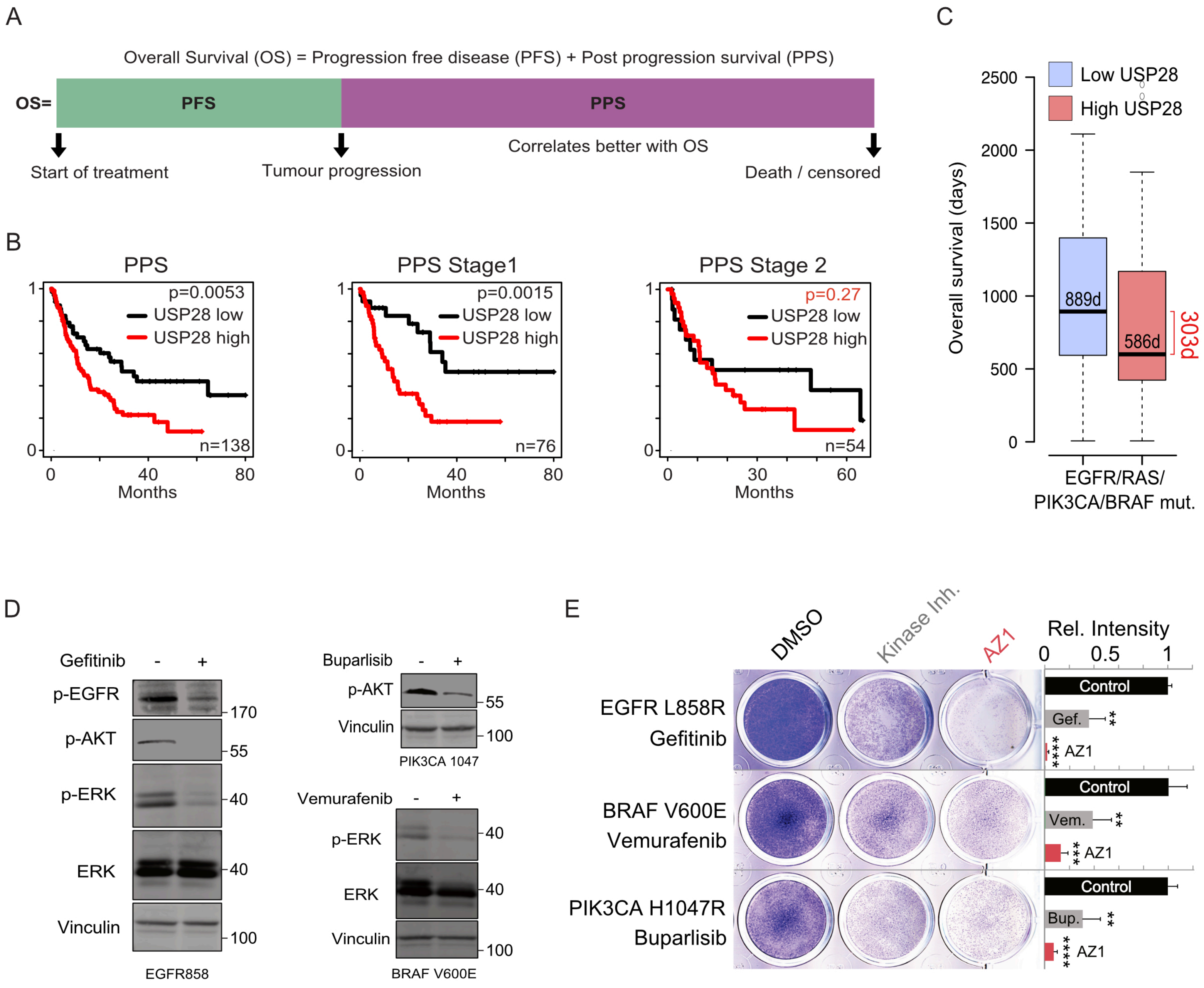


Prieto-Garcia et al. Figure S7
